# A novel cost-benefit decision-making task involving cued punishment; effects of sex and psychostimulant administration

**DOI:** 10.1101/2025.07.15.664953

**Authors:** Mojdeh Faraji, Jennifer L. Bizon, Barry Setlow

## Abstract

Chronic substance use is associated with alterations in multiple forms of cost-benefit decision making, which may prolong and exacerbate continued use. Cues that predict reward can cause substantial shifts in a variety of reward-directed behavior, including decision making. In contrast, how decision making is modulated by cues predictive of punishment is much less well understood. To begin to address these issues, male and female Long-Evans rats were tested in a novel decision-making task in which they chose between a small, “safe” reward and a large reward that was punished by a mild footshock when it was preceded by a probabilistically delivered cue prior to the choice. Rats of both sexes were sensitive to the cue, preferring the large reward in the absence of the cue but the small reward in the presence of the cue. Acute systemic amphetamine reduced choice of the large reward and diminished the efficacy of the cue in guiding choice behavior. Chronic cocaine led to divergent patterns of cue insensitivity in males and females; males increased choice of the large reward on cued trials, whereas females increased avoidance of the large reward on uncued trials. Similar to acute amphetamine, acute systemic administration of the D2/3 dopamine receptor agonist bromocriptine reduced preference for the large reward across all groups. These findings highlight the contributions of punishment cues to decision making, as well as the importance of sex as a biological variable in investigating cognitive alterations caused by chronic substance use.

## 1. Introduction

It is estimated that more than 41 million people in the United States alone are living with a substance use disorder[1]. Substance use disorders are associated with cognitive impairments across multiple domains, including executive functions and decision making [2–11]. In particular, chronic substance use is frequently accompanied by alterations in several forms of cost-benefit decision making, such as elevated preference for risky vs. safe choices [12–14]. Dysregulated cost-benefit decision making in individuals with a history of substance use can lead to devastating consequences (e.g. overdose, intoxicated driving, and disease transmission through intravenous injection and unsafe sexual activity). Altered evaluation of adverse outcomes may promote continued substance use by promoting preference for the risky rewards of substance use over the safer rewards of abstinence. In addition, as elevated risk taking can persist even long into abstinence, it may increase the likelihood of relapse. Such behavior is evident in laboratory tasks such as the Iowa Gambling Task, Cambridge Gamble Task, and Balloon Analogue Risk Task, in which users of cocaine and other stimulants make more risky choices, resulting in overall worse task performance [5, 15–20].

Elevated risk taking is one of the prominent manifestations of cognitive impairments in substance use disorders. Rodent models have been long used to study risky decision making by employing operant chamber assays that evaluate cost-benefit decision making in conflict situations. Typically, the task designs in these assays provide a no-contingency option accompanied by a small reward and a larger reward accompanied by a “cost” (e.g. reward omission, footshock, aversive taste). In previous work from our lab, we developed a rat model of risky decision making (the “risky decision-making task”; RDT) [21, 22] that mirrors numerous features of human behavior relevant to risk taking in the context of substance use. In that task, rats make discrete choices between a small, “safe” food reward and a large food reward that is accompanied by varying probabilities of footshock punishment [21]. Rats with a history of cocaine self-administration show long-lasting increases in preference for large, risky over small, safe rewards in this task [23, 24], mimicking patterns of behavior evident in people with a history of cocaine use [12, 25, 26].

Much of the existing research on the effects of substance use on motivated behavior has focused on drug-induced changes in the value of rewards rather than the associated costs. As such, it is not clear to what extent alterations in risk-based decision making in both humans and rodent models in the context of substance use are due to impairments in risk evaluation or attenuated sensitivity to negative outcomes more broadly. In humans, stimulant drug use is associated with blunted neural responses to monetary loss in gambling tasks [27, 28]. In rodents, prolonged cocaine exposure has been shown to diminish the effects of punishment on drug-seeking behavior [29], and attenuate motivation to avoid aversive outcomes [30], offering evidence for reduced sensitivity to punishment. On the other hand, drug-induced punishment resistance has been reported to be positively sensitive to the uncertainties accompanied by the punished choice [31], and uncertain cocaine-paired cues were shown to increase nucleus accumbens dopamine signals significantly more than certain cocaine-paired cues [32]. Here we extend our previous work with the RDT by examining whether heightened preference for large, risky rewards in the RDT following chronic cocaine is due to discounting the risk of punishment or desensitization to the punishment itself. Addressing this issue could help in designing therapeutic approaches as one (discounted risk) might assign greater weight to the role of environmental factors and proximity to risky environments, and the other (desensitization to harm) might assign greater weight to disrupted neurocognitive factors in treatment approaches.

To address these questions, we used a novel variant of the RDT that disentangles risk from the aversive outcome. The “cued punishment decision-making task” (CPDT) eliminates the risk component in the RDT by using a blinking light cue that signals that punishment (footshock) will accompany choice of a large reward. In this task, delivery of punishment is cued in advance, prior to the choice phase when both small and large reward levers become available. On cued trials, choice of the large reward is accompanied by a mild footshock and on uncued trials, choice of the large reward is unpunished. In other words, by using a deterministic cued punishment, the CPDT determines whether environmental cues are effective in shifting choice behavior to avoid adverse outcomes. We then investigated how these processes are affected by acute and chronic stimulant administration (amphetamine and cocaine, respectively). Finally, we evaluated how acute administration of the D2/3 dopamine receptor agonist bromocriptine (which reduces preference for large, probabilistically-punished rewards in the RDT [33, 34]), affected decision making in the context of deterministic punishment.

Prior research shows that repeated cocaine causes a lasting reduction in motivation to avoid aversive outcomes [30]. Hence, our hypothesis was that drug-naïve rats would use information from the punishment cue to avoid footshock on cued trials by choosing the small reward, while maximizing their food intake on uncued trials by choosing the large reward. Based on our and others’ prior work showing that females are more sensitive to punishment [27, 35], we anticipated that females would be more attentive to the punishment cue and make fewer choices of the large reward on cued trials compared to males. Finally, we expected that in both sexes, acute exposure to dopaminergic agonists would reduce choice of the large reward on cued trials, whereas chronic cocaine exposure, similar to our previous results in the RDT [24], would produce a lasting increase in choice of the large reward on cued trials.

## 2. Materials and Methods

### 2.1. Subjects

Male and female Long-Evans rats (N=30, bred in-house from founders originating at Charles River Laboratories, Raleigh, NC) were housed in sex-matched pairs after weaning until 50 days of age (P50), at which point they were housed individually. Rats were tested in two cohorts. Those in Experiment 1 (n=6 male, n=6 female) were food restricted beginning at P60, and started behavioral testing once they reached 85% of their free-feeding weight. Rats in Experiment 2 (n=9 male, n=9 female) received cocaine or saline injections (see Experimental Design below) beginning at P60, and were food restricted to 85% of their free-feeding weight beginning at P95, followed by behavioral testing. During periods of food restriction, rats’ target weights were adjusted upward by 5 g/week to account for growth until 6 months old. Rats were kept on a 12- hour light/dark cycle (lights on at 0700), maintained at a consistent temperature of 25°C, fed on Teklad irradiated global 19% protein chow (2919) and had access to water *ad libitum*. Animal procedures were conducted in accordance with the University of Florida Institutional Animal Care and Use Committee and followed guidelines of the National Institutes of Health.

### 2.2. Apparatus

Behavioral testing was conducted in standard operant chambers (Coulbourn Instruments). Chambers were contained in sound-attenuating cubicles, and were computer-controlled through Graphic State 4.0 software (Coulbourn Instruments). Locomotor activity was monitored via infrared motion detectors installed on the chamber ceiling. Each operant chamber was equipped with a food trough containing a photobeam sensor to detect nosepokes into the trough, two retractable levers (one on each side of the food trough), a feeder installed on the outside wall of the chamber and connected to the food trough to deliver 45 mg purified ingredient rodent food pellets (Test Diet; 1811155 5TUM), and a stainless steel floor grate connected to a shock generator that delivered scrambled footshocks. Each sound-attenuating cubicle included a house light mounted on the rear wall (outside of the operant chamber).

### 2.3. Behavioral procedures

#### 2.3.1. Cued Punishment Decision-making Task

Prior to testing in the cued punishment decision-making task (CPDT), rats were trained on a sequence of shaping protocols to learn how to retrieve food from the food trough, nosepoke in the food trough to initiate a trial, and press the levers to obtain food. Shaping procedures are described in detail in [24]. Upon completion of shaping, rats proceeded to the CPDT itself. The structure of the CPDT is based on the risky decision-making task (RDT) used previously in our lab and elsewhere [21, 22]. Unlike the RDT, in which choice of the large reward is followed by a mild shock in a probabilistic manner, in the CPDT, shock is fully predicted by a light cue that precedes each choice. On each trial in this task, rats had the opportunity to press one of two levers (Figure 1). Trials began with illumination of the food trough. A nosepoke into the trough extinguished the trough light and triggered a 3-s period during which a flashing (1 Hz) light cue indicating punishment upon selection of the large reward lever was delivered on a probabilistic schedule. Failure to nosepoke in the trough within 10 s caused the trough light to extinguish, and the trial was counted as an omission. The cue light period was followed by extension of one (forced-choice trials) or both (free-choice trials) levers. A press on one of the levers led to delivery of 1 food pellet (small reward). A press on the other lever led to delivery of 2 food pellets (large reward), which was accompanied by a mild footshock (1.0 s in duration) on trials on which the cue light preceded lever extension. Thus, the presentation of the cue light indicated that choice of the large reward would be punished on that trial. Lever presses were followed by retraction of the lever(s), illumination of the food trough light, and delivery of food pellets. The food trough light was extinguished upon retrieval of the pellets or after 10 s, whichever came first. Trials were separated by an intertrial interval (ITI) in which the house light was extinguished.

**Figure 1.**
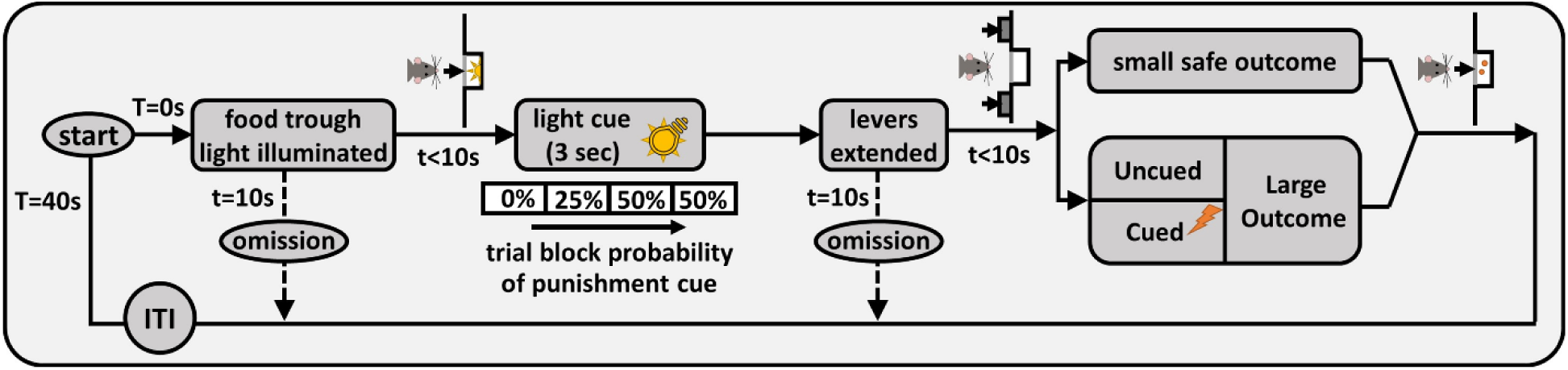
Cued punishment decision-making task (CPDT) schematic. Each trial was initiated by a nosepoke into the illuminated food trough, which was followed by a probabilistic delivery of a cue light (3 s). Levers were extended when the cue light was extinguished, and pressing the levers led to delivery of food pellet(s). If the large reward lever was chosen on a cued trial, delivery of this reward was accompanied by footshock delivery. There were four blocks of trials in the CPDT, and the probability of cue light delivery increased as blocks proceeded. “T” denotes the total time spent in a trial and “t” denotes the time spent in each state of the trial.

Each session was comprised of 4 blocks of trials with different probabilities of cue light delivery (0, 25, 50, 50%). Each block of trials began with 8 forced-choice trials (4 for each lever) in which only one lever was extended. The goal of these trials was to provide a reminder regarding the probability of cue light delivery (and consequent shock if the large reward were selected) in effect in that block. The forced-choice trials were followed by 10 (blocks 1 and 2) or 15 (blocks 3 and 4) free-choice trials in which both levers were extended. Sessions in the CPDT were 55 minutes in duration and consisted of 82 trials, each 40 s long. The left/right positions of the small and large reward levers were randomized across rats, but remained consistent for each rat over the course of the task. Shock intensities were 300𝜇A for females and 400 𝜇A for males in Experiment 1, and 375𝜇A for all rats in Experiment 2.

### 2.4. Experimental procedures

#### 2.4.1. Experiment 1

Rats (n=6 male, n=6 female) were trained on the CPDT until stable performance emerged (see Data Analysis section for description of performance stability). Beginning at 7 months old, the effects of systemic administration of amphetamine on task performance were evaluated, using a regimen shown previously to reduce choice of large, risky rewards in the RDT [35, 36]. D-amphetamine (NIDA Drug Supply Program, 0, 0.3, 1.0, 1.5 mg/kg, intraperitoneal (IP), 0.9% saline vehicle, 1.0 ml/kg) was administered using a randomized, within-subjects design such that each rat received each dose, with a 48-h washout period between successive doses. Injections took place 10 min prior to testing in the CPDT.

#### 2.4.2. Experiment 2

In Experiment 2, effects of chronic cocaine exposure on performance in the CPDT were evaluated. Rats were divided into two groups (control and cocaine). Beginning at P60, the cocaine group received daily injections of cocaine (n=4 male, n=5 female) or 0.9% saline vehicle (n=5 male, n=4 female) for 14 consecutive days. The doses of cocaine HCl (NIDA Drug Supply Program, IP) were as follows: 15 mg/kg for 3 days, 20 mg/kg for 2 days, 25 mg/kg for 2 days, and 30 mg/kg for 7 days. This escalating dose regimen is based on cocaine regimens shown previously to produce lasting alterations in several forms of cost-benefit decision making [24, 37]. Following completion of the injection regimen, rats remained undisturbed in their home cages for 3 weeks, followed by food restriction and training in the CPDT until stable performance emerged.

Beginning at 6 months old the effects of systemic administration of the D2/3 dopamine receptor agonist bromocriptine on task performance were evaluated, using a regimen shown previously to reduce choice of large, risky rewards in the RDT [33, 34]. Bromocriptine mesylate (Tocris Bioscience, 0, 1.0, 3.0, 5.0 mg/kg, IP, 1.0 ml/kg, in a 1:1 DMSO and 0.9% saline vehicle) was administered using a randomized, within-subjects design such that each rat received each dose, within a 48-h washout period between successive doses. Injections took place 40 min prior to behavioral testing.

### 2.5. Data analysis

Data were collected and processed using custom protocols and analysis templates in Graphic State 4.0. Statistical analyses were conducted in SPSS and graphs created using GraphPad Prism 9. To evaluate stable performance in the CPDT, two-factor repeated measures analyses of variance (ANOVA) with session and trial block as within-subjects factors were conducted on choice data across 3 consecutive sessions within each group (Experiment 1, males and females; Experiment 2, males and females in control and cocaine groups). Stability was defined as the absence of a significant main effect of Session or Session x Cue Probability interaction (p>0.05).

The first measure of interest in the CPDT was the percentage of choices of the large reward in each block (out of the total choices made in each block). In Experiment 1, the percentage of choices of the large reward at baseline was evaluated using a two-factor repeated-measures ANOVA, with Sex as a between-subjects factor and Cue Probability as a within-subjects factor. The effects of amphetamine administration on the percentage of choices of the large reward were evaluated using a three-factor repeated-measures ANOVA, with Sex as a between-subjects factor and Amphetamine and Cue Probability as within-subjects factors.

To examine the immediate effects of the punishment cue on decision making, rats’ choice distributions (total number of cued small, cued large, uncued small, and uncued large choices) in trial blocks 3 and 4 were analyzed during free-choice trials. These analyses focused on blocks 3 and 4 as the probability of cue presentation was 50%, resulting in roughly equivalent preference for the small and large reward levers at baseline. On free-choice trials, both levers are presented simultaneously, and choice of either lever is a dependent variable. Therefore, Lever identity was not included as an independent ANOVA factor in this analysis, and large reward choices (cued and uncued) were analyzed separately from small reward choices (cued and uncued). Results were compared between males and females at baseline using a two-factor repeated measures ANOVA, with Sex as a between-subjects factor and Cue as a within-subjects factor on data averaged across stable sessions. The effects of the punishment cue on decision making following amphetamine administration were evaluated using a three-factor repeated measures ANOVA, with Sex as a between-subjects factor and Amphetamine and Cue as within-subjects factors.

Ancillary measures, including locomotor activity, shock reactivity (locomotor activity during shock delivery), omissions, and total rewards earned during free-choice trials were analyzed using Welch’s t-tests, with Sex as the group factor. Latencies to press levers on free-choice trials during blocks 3 and 4 were analyzed using a three-factor repeated measure ANOVA, with Sex as a between-subjects factor, and Lever identity and Cue as within-subjects factors. Ancillary measures following amphetamine administration were analyzed using a two-factor repeated-measure ANOVA, with Sex as a between-subjects factor and Amphetamine as a within-subjects factor. Latencies to press levers in free choice trials in blocks 3 and 4 were analyzed using a multi-factor repeated measure ANOVA, with Sex as a between-subjects factor, and Lever identity, Cue, and Amphetamine as within-subjects factors.

In Experiment 2, the percentage of choices of the large reward at baseline was evaluated using a three-factor repeated measures ANOVA, with Sex and Cocaine group as between-subjects factors and Cue Probability as a within-subjects factor. The effects of bromocriptine on the percentage of choices of the large reward were evaluated using a multi-factor repeated measures ANOVA, with Sex and Cocaine group as between-subjects factors and Bromocriptine and Cue Probability as within-subjects factors. The choice distribution results (during blocks 3 and 4 only) were compared between groups using a three-factor repeated-measures ANOVA, with Sex and Cocaine group as between-subjects factor and Cue as a within-subjects factor on data averaged across stable sessions. Similar to Experiment 1, Lever identity was not included as an independent ANOVA factor in this analysis, and large reward choices (cued and uncued) were analyzed separately from small reward choices (cued and uncued). The same analysis following bromocriptine administration was conducted using a multi-factor repeated measures ANOVA, with Sex and Cocaine group as between-subjects factors and Bromocriptine and Cue as within-subjects factors.

Ancillary measures including locomotor activity, shock reactivity (locomotor activity during shock delivery), and omissions during free choice trials were analyzed using a two-factor repeated measures ANOVA, with Sex and Cocaine group as between-subjects factors. Latencies to press levers in free-choice trials in blocks 3 and 4 were analyzed using a multi-factor repeated measures ANOVA, with Sex and Cocaine group as between-subjects factors, and Lever identity and Cue as within-subjects factors. Ancillary measures following bromocriptine administration were analyzed using a three-factor repeated-measure ANOVA, with Sex and Cocaine group as between-subjects factors and Bromocriptine as a within-subjects factor.

Latencies to press levers in free choice trials were analyzed using a multi-factor repeated measures ANOVA, with Sex and Cocaine group as between-subjects factors and Lever identity, Cue, and Bromocriptine as within-subjects factors. For all analyses, the Greenhouse-Geisser correction was used to account for violations of sphericity in ANOVAs, and p values less than or equal to .05 were considered significant.

## 3. Results

### 3.1. Experiment 1

#### 3.1.1. Choice of large reward

Rats were tested on the CPDT until stable performance emerged (about 40 sessions). To achieve comparable choice preference between males and females before administration of amphetamine, different shock intensities (females, 300 𝜇A; males, 400 𝜇A) were used in the two sexes in this experiment [35]. At these shock intensities (Figure 2a), a two-factor repeated measures ANOVA (Sex x Cue Probability) on % choice of large reward revealed a main effect of Cue Probability (F(1.67,16.71)=24.07, p<0.01) such that choice of the large reward decreased as the probability of punishment cue increased, but choice behavior did not differ between males and females (Sex, F(1,10)=0.84, p=0.38), nor was there an interaction between Sex and Cue Probability (F(2,20)=0.02, p=0.98). Note that the percent choice of the large reward includes choice of the large reward on both cued and uncued (punished and unpunished) trials.

**Figure 2.**
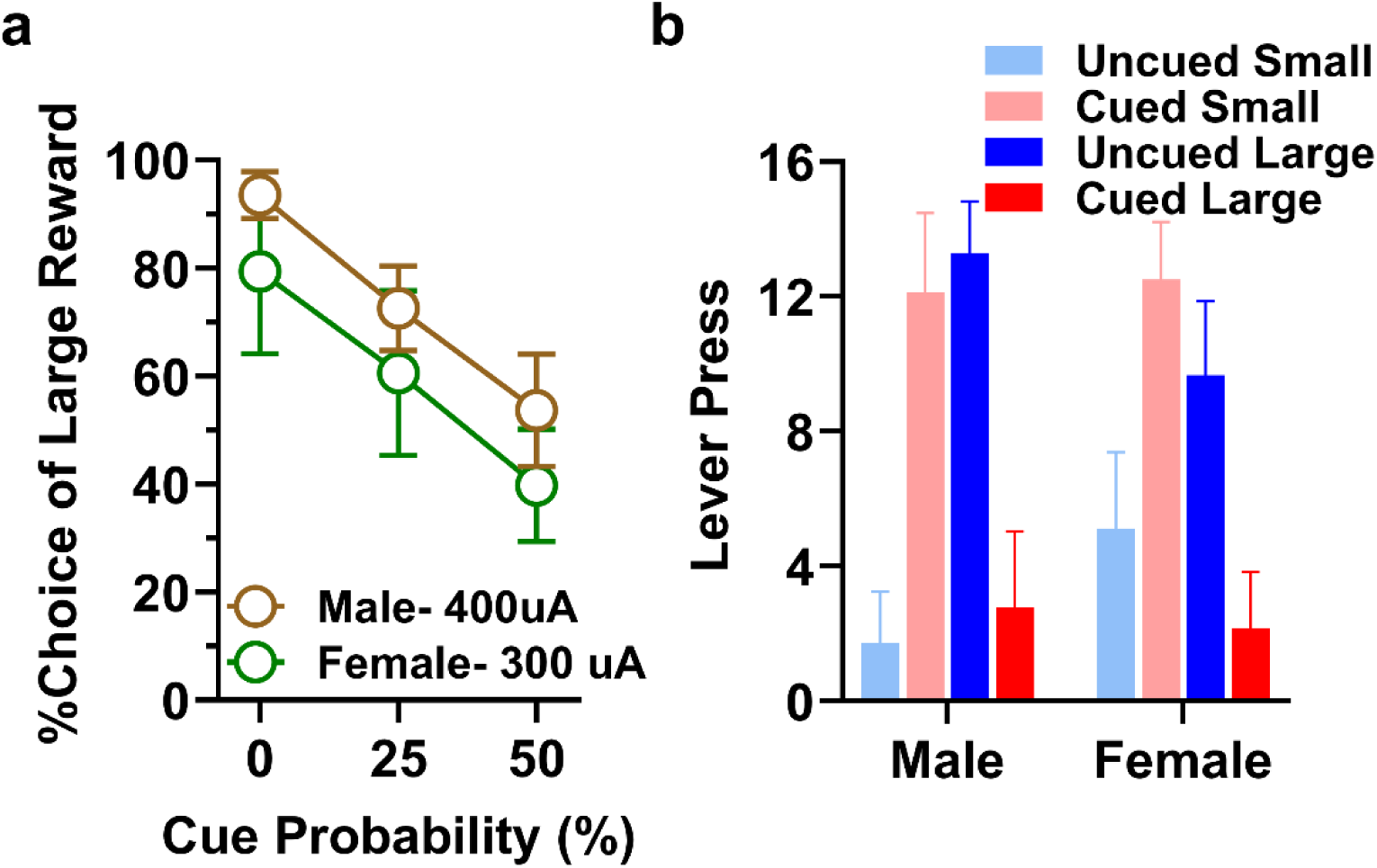
Experiment 1, Choice behavior in the CPDT. **a)** %Choice of the large reward decreased as the probability of the punishment cue increased, but there was no main or interaction effect of sex. **b)** The presence of the punishment cue decreased the number of large reward choices (p<0.01), but results did not differ between males and females. Data are represented as means ± SEM.

#### 3.1.2. Choice distribution at 50% probability of punishment cue

To further evaluate the effects of the punishment cue on choice behavior, subsequent analyses focused on trial blocks in which the probabilities of cued (punished) and uncued (unpunished) trials were equal (50% cue probability). There were four possible combinations of trial types and outcomes. For uncued (unpunished) trials, rats could choose either the small or large reward and earn the corresponding number of food pellets without further consequences. On these trials, choice of the large reward lever could thus be considered “optimal” by maximizing the amount of reward earned, and choice of the small reward lever could be considered suboptimal. For cued (punished) trials, choice of the large reward lever led to large reward delivery accompanied by a footshock, and choice of the small reward lever led to the small reward delivery without further consequences. Hence, on these trials, choice of the large reward represents a “punishment-insensitive” choice, and the small reward lever could be considered the “safe” choice by avoiding the punishment.

To evaluate how rats incorporate information about the punishment cue into their choice behavior, we compared the choice distribution between males and females during the 50% cue probability blocks (Figure 2b). A two-factor repeated measures ANOVA (Sex x Cue) on the number of large reward choices revealed a main effect of Cue (such that the large reward was chosen more on uncued compared to cued trials; Cue, F(1,10)=30.28, p<0.01) but no effect of Sex or Sex x Cue interaction (Sex, F(1,10)=0.92, p=0.36; Sex × Cue, F(1,10)=0.84, p=0.38). The same analysis conducted on small reward choices revealed comparable results (Cue, F(1,10)=27.01, p<0.01; Sex, F(1,10)=0.70, p=0.42; Sex × Cue, F(1,10)=0.77, p=0.40). These findings indicate that the appearance of the cue prior to the choice shifted rats’ preference from the large to the small reward in both males and females (greater number of large reward choices on uncued trials and greater number of small reward choices on cued trials).

Latencies to press levers on free-choice trials at equal probability of cue appearance (50%) were analyzed using a three-factor repeated measures ANOVA (Sex x Cue x Lever). There was an interaction between Cue and Lever (F(1,20.86)=10.91, p<0.01) such that latencies to press the large reward lever were longer on cued compared to uncued trials (possibly reflecting longer deliberation times in anticipation of punishment), whereas latencies to press the small reward lever were shorter on cued compared to uncued trials. There were no main effects of Sex, Cue, or Lever, nor were there any other interactions among the factors on lever press latencies (Tables 1 and 2).

**Table 1.**
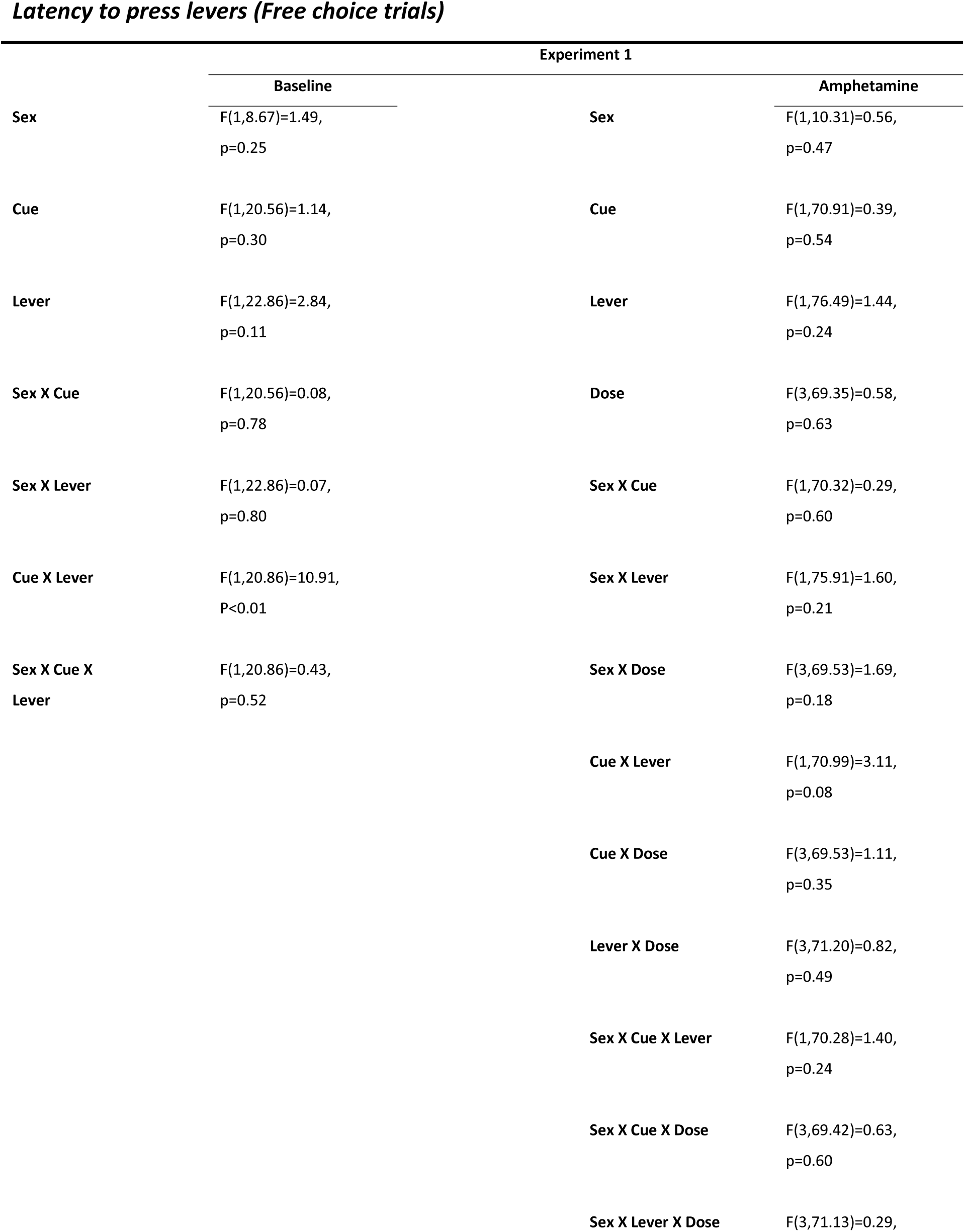
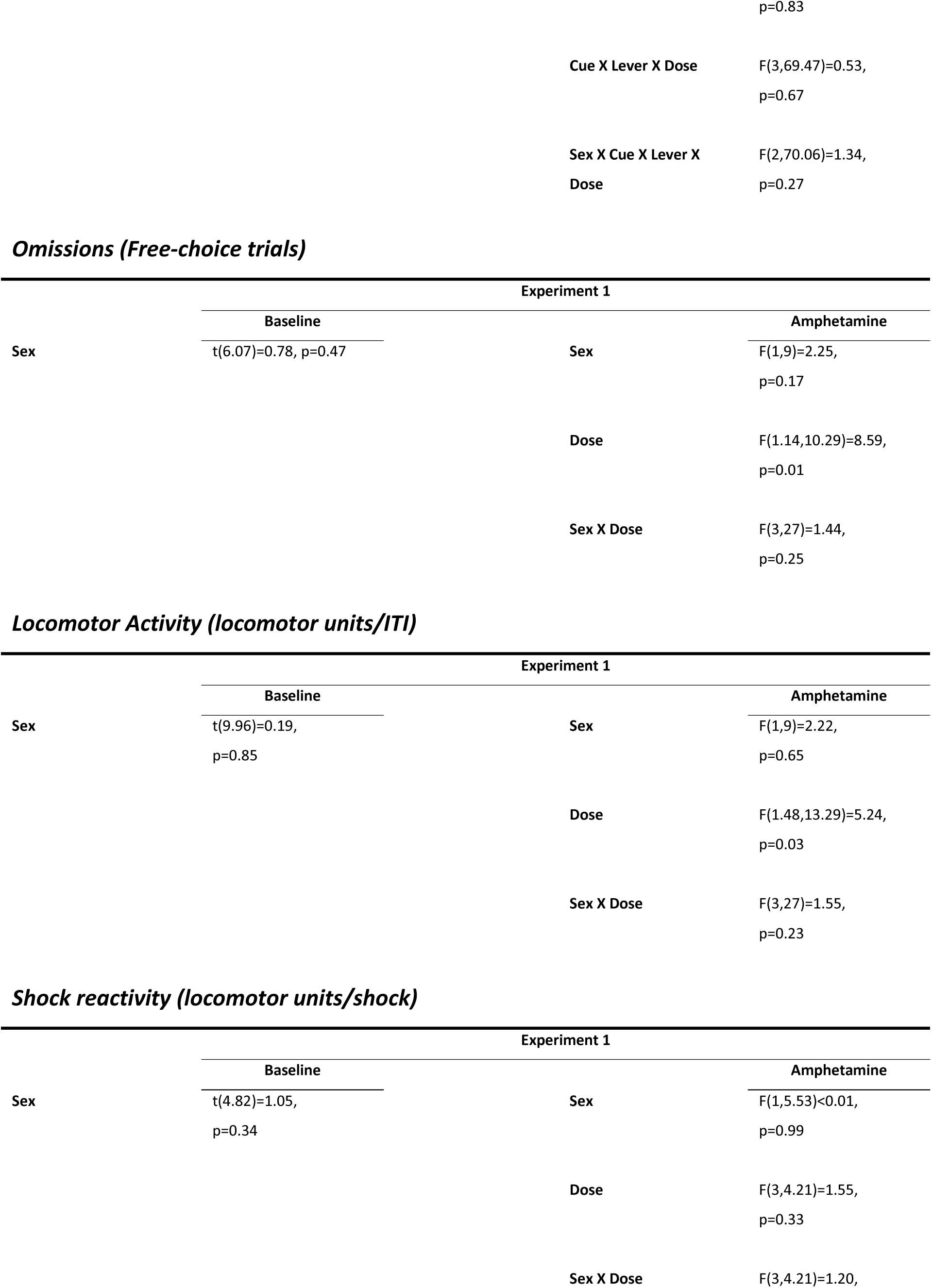
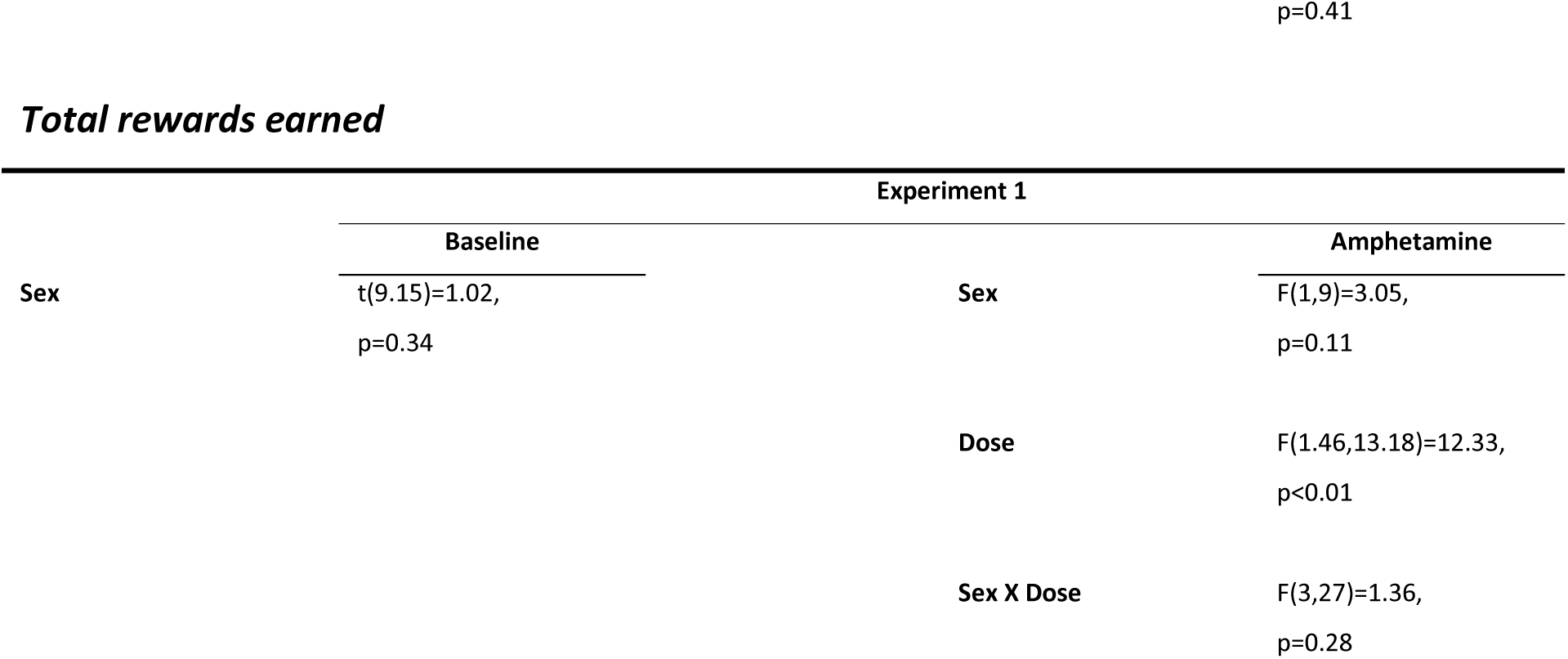
Ancillary measures (statistical results), experiment 1.

**Table 2.**
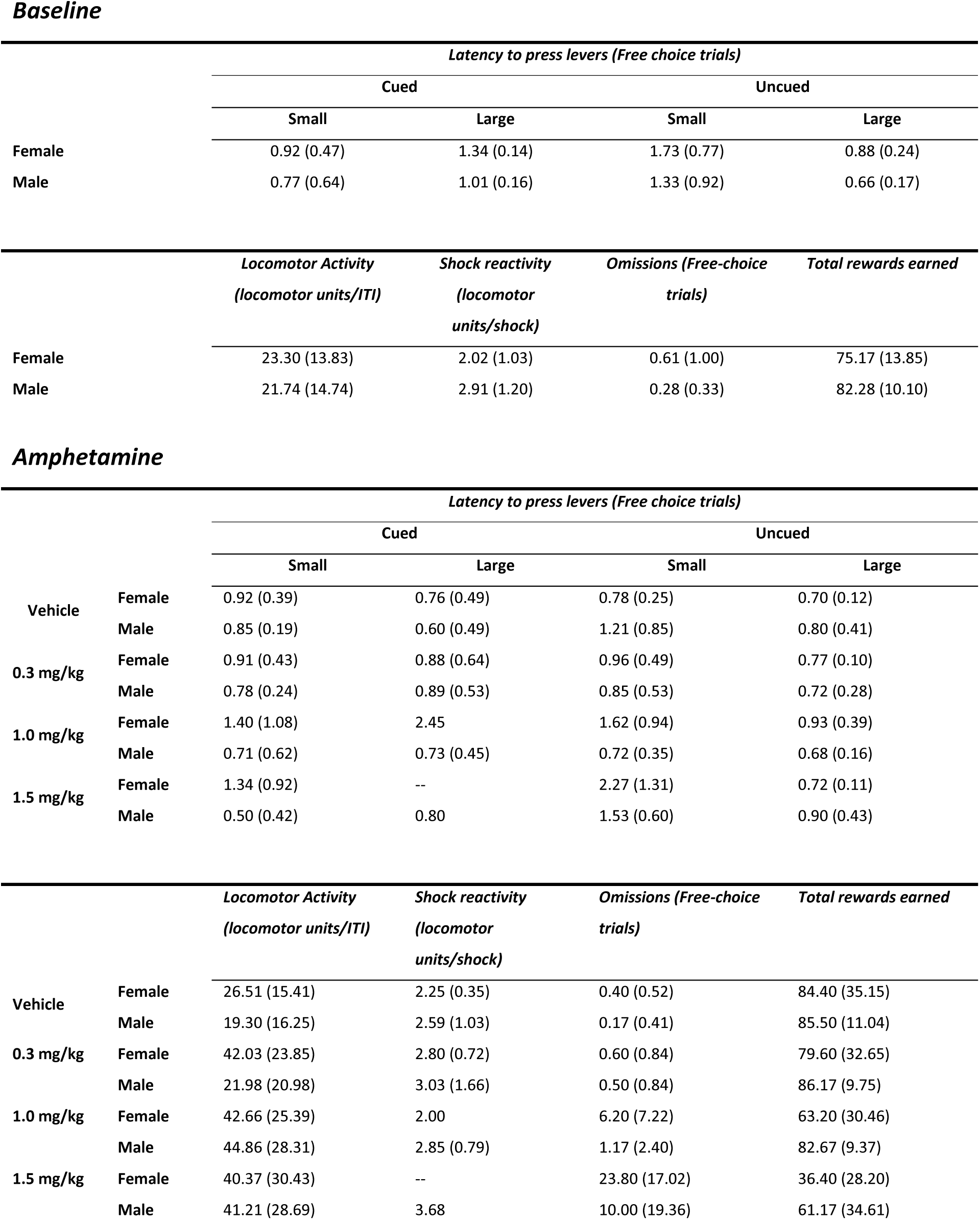
Ancillary measures (mean ± standard error of the mean), experiment 1.

Comparison of locomotor activity in the CPDT between males and females using a Welch’s t-test revealed no sex differences during either intertrial intervals (t(9.96)=0.19, p=0.85), or during shock deliveries in the task (t(4.82)=1.05, p=0.34). Analysis of the number of free-choice trial omissions (Welch’s t-test) also revealed no effects of Sex (t(6.07)=0.78, p=0.47), nor was there any difference between sexes in total number of rewards earned (Welch’s t-test, t(9.15)=1.02, p=0.34; Tables 1 and 2).

#### 3.1.3. Amphetamine

Performance on the CPDT was tested following acute systemic administration of amphetamine (Figure 3). A three-factor repeated measures ANOVA (Sex x Amphetamine x Cue Probability) on % choice of the large reward revealed the expected main effect of Cue Probability (F(2,93.44)=34.10, p<0.01) to reduce choice of the large reward, and a main effect of Amphetamine (F(3,93.50)=13.10, p<0.01) such that choice of the large reward decreased following administration of amphetamine in a dose-dependent manner (Figure 3a, c). Dunnett’s post-hoc comparisons revealed that compared to vehicle, amphetamine at 1.5 mg/kg significantly reduced choice of the large reward (p<0.01). Neither a main effect of Sex (F(1,9.26)=3.07, p=0.11) nor any interactions concerning Sex or Amphetamine were observed (Sex x Amphetamine, F(3,93.50)=2.07, p=0.11; Sex x Cue Probability, F(2,93.44)=0.41, p=0.67; Amphetamine x Cue Probability, F(6,93.41)=0.50, p=0.81; Sex x Amphetamine x Cue Probability, F(6,93.41)=0.39, p=0.89).

**Figure 3.**
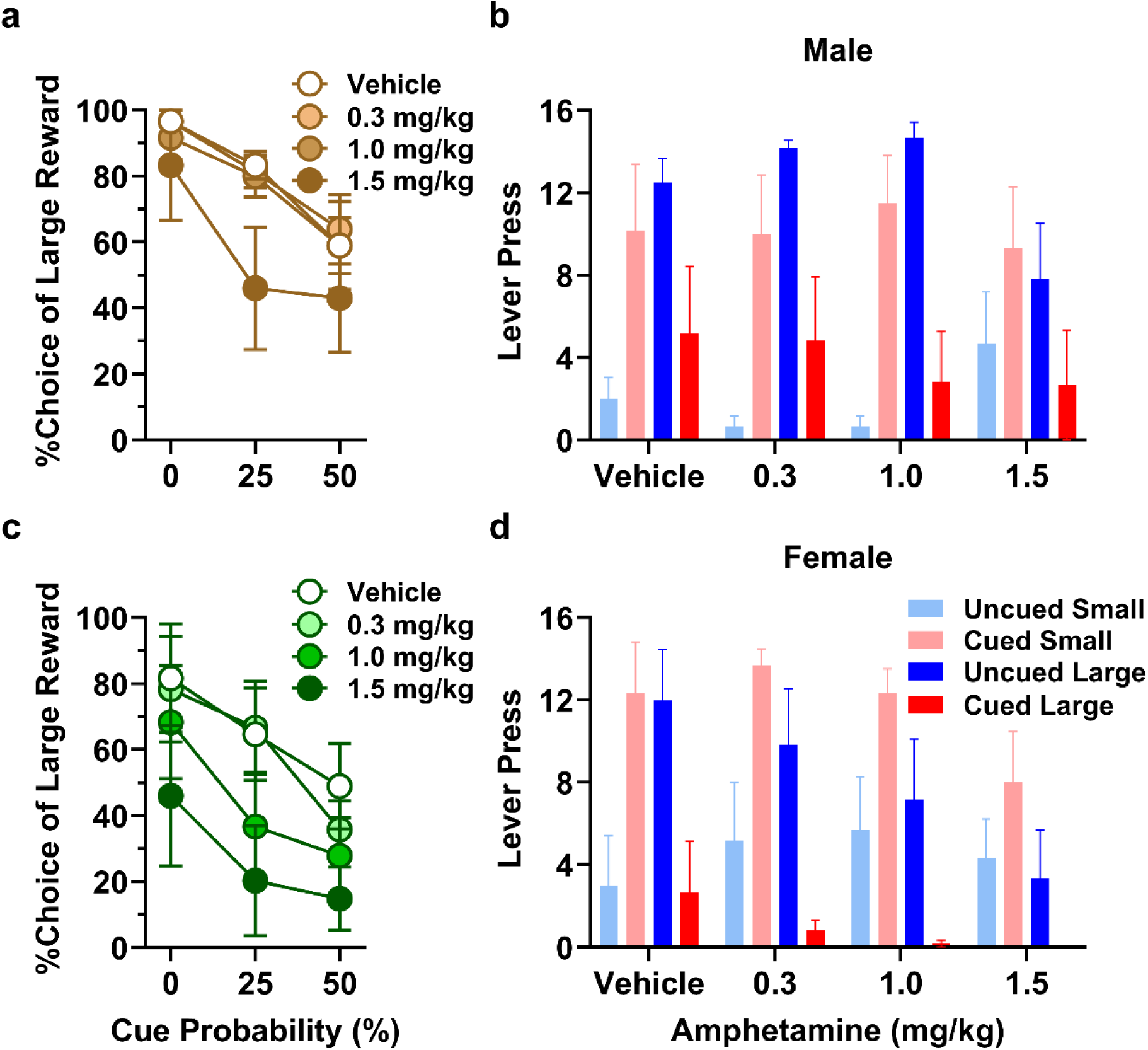
Experiment 1, Effects of acute amphetamine on choice behavior in the CPDT. **a, c)** %Choice of the large reward decreased as the probability of the punishment cue increased (p<0.01), and with administration of amphetamine (p<0.01), but the results did not differ between males (a) and females (c). **b, d)** Both the cue (p<0.01) and amphetamine (p<0.01) decreased the number of large reward choices, but results did not differ between males (b) and females (d). Data are represented as means ± SEM.

Analysis of the effect of punishment cue (three-factor repeated measures ANOVA, Sex x Amphetamine x Cue) on rats’ choice of the large reward at 50% cue probability revealed a main effect of Cue (F(1,9)=28.74, p<0.01) and a main effect of Amphetamine (F(1.56,14.08)=7.40, p<0.01) such that rats chose the large reward less frequently on cued vs. uncued trials, and that amphetamine reduced the number of large reward lever presses (Figure 3b, d). There was, however, no significant interaction between Amphetamine and Cue (F(1.44,12.98)=3.41, p=0.08). A Dunnett’s post-hoc multiple comparison test revealed that choice of the large reward following amphetamine was significantly reduced at the 1.5 mg/kg dose compared to vehicle (p<0.01). No main effect or interaction involving Sex was evident (Sex, F(1,9)=1.99, p=0.19; Amphetamine x Sex: F(3,27)=1.23, p=0.32; Cue x Sex, F(1,9)<0.01, p=0.96; Amphetamine x Cue x Sex, F(3,27)=1.22, p=0.32). A similar analysis of rats’ choice of the small reward at 50% cue probability revealed a main effect of Cue (F(1,9)=29.61, p<0.01) such that rats chose the small reward more frequently on cued vs. uncued trials. There was an interaction between Amphetamine and Cue (F(2.04,18.33)=3.50, p=0.05), and a Bonferroni’s post-hoc multiple comparisons test revealed that choice of the small reward was significantly more frequent on cued trials at vehicle and following amphetamine administration at 0.3 and 1.0 mg/kg (p<0.01), but following amphetamine administration at 1.5 mg/kg the effect of cue on choice of small reward was diminished (p=0.16). There was no main effect of Amphetamine or Sex, nor any interactions involving Sex, on choice of the small reward (Amphetamine, F(1.64,14.74)=0.59, p=0.54; Sex, F(1,9)=0.29, p=0.60; Amphetamine x Sex: F(3,27)=1.66, p=0.20; Cue x Sex, F(1,9)<0.01, p=0.95; Amphetamine x Cue x Sex, F(3,27)=0.77, p=0.52).

Latencies to press levers on free-choice trials during blocks 3 and 4 were analyzed using a multi- factor repeated measures ANOVA (Sex x Cue x Lever x Amphetamine). Amphetamine increased latencies (F(3,97.48)=3.95, p=0.01) and females showed longer latencies compared to males (F(1,11.42)=7.53, P=0.02). There was also an interaction between Amphetamine and Sex (F(3,97.23)=6.03, p<0.01) such that amphetamine increased lever press latencies in females but decreased latencies in males, and an interaction between Cue and Lever (F(1,97.16)=6.69, p=0.01), such that latencies to press the large reward lever were longer on cued compared to uncued trials, whereas latencies to press the small reward lever were shorter on cued compared to uncued trials. There was no main effect of Cue or any other interaction between the factors on latency to press levers (Tables 1 and 2).

A two-factor repeated measures ANOVA (Sex x Amphetamine) was used to evaluate locomotor activity during the CPDT. Amphetamine increased locomotor activity during the ITI (F(1.48,13.29)=5.24, p=0.03), but there was no main effect of or interaction concerning Sex (Sex, F(1,9)=2.22, p=0.65; Amphetamine x Sex, F(3,27)=1.55, p=0.23). Post-hoc Dunnett’s multiple comparisons tests revealed that the increase in locomotor activity during the ITI was statistically significant at the 1 and 1.5 mg/kg doses. Locomotor activity during shock deliveries in the task was not affected by amphetamine, nor did it differ between males and females (Tables 1 and 2).

A two-factor repeated measures ANOVA (Amphetamine x Sex) conducted on the number of omitted trials revealed that Amphetamine increased trial omissions (F(1.14,10.29)=8.59, p=0.01), but there was no main effect or interaction concerning Sex (Sex, F(1,9)=2.25, p=0.17; Amphetamine x Sex, F(3,27)=1.44, p=0.25). A Dunnett’s post-hoc multiple comparisons test revealed that the number of omitted trials following amphetamine administration was significantly increased at the 1.5 mg/kg dose compared to vehicle (p=0.04; Tables 1 and 2).

Analysis of total number of rewards earned (two-factor repeated measures ANOVA, Sex x Amphetamine) during the CPDT revealed a main effect of Amphetamine (F(1.46,13.18)=12.33, p<0.01) such that amphetamine reduced the number of rewards earned. There was, however, no main effect of Sex (F(1,9)=3.05, p=0.11), nor was there an interaction between Sex and Amphetamine (F(3,27)=1.36, p=0.28). A Dunnett’s post-hoc multiple comparisons test revealed that compared to vehicle, the 1.5 mg/kg dose significantly reduced the total rewards earned (p=0.01; Tables 1 and 2).

### 3.2. Experiment 2

#### 3.2.1. Choice of large reward

Following a 14-day regimen of cocaine or saline vehicle injections, rats (control male, n=5; cocaine male, n=4; control female, n=4; cocaine female, n=5) were tested on the CPDT until stable performance emerged (roughly 25 sessions; Figure 4a). A three-factor repeated measures ANOVA (Cocaine x Sex x Cue Probability) on % choice of the large reward revealed a main effect of Cue Probability (F(1.55,21.69)=51.54, p<0.01) such that preference for the large reward declined as probability of punishment cue increased. Male rats chose the large reward more frequently than females (Sex, F(1,14)=8.96, p<0.01), and there was an interaction between Sex and Cocaine (F(1,14)=4.91, p=0.04) such that in males, cocaine increased choice of the large reward compared to controls, but decreased the same measure in females. The main effect of Cocaine, however, was not significant (F(1,14)=0.13, p=0.73). No other interaction concerning Sex or Cocaine was significant (Cue Probability × Sex, F(2,28)=1.19, p=0.32; Cue Probability × Cocaine, F(2,28)=0.41, p=0.67; Cue Probability × Cocaine × Sex, F(2,28)=0.97, p=0.39).

**Figure 4.**
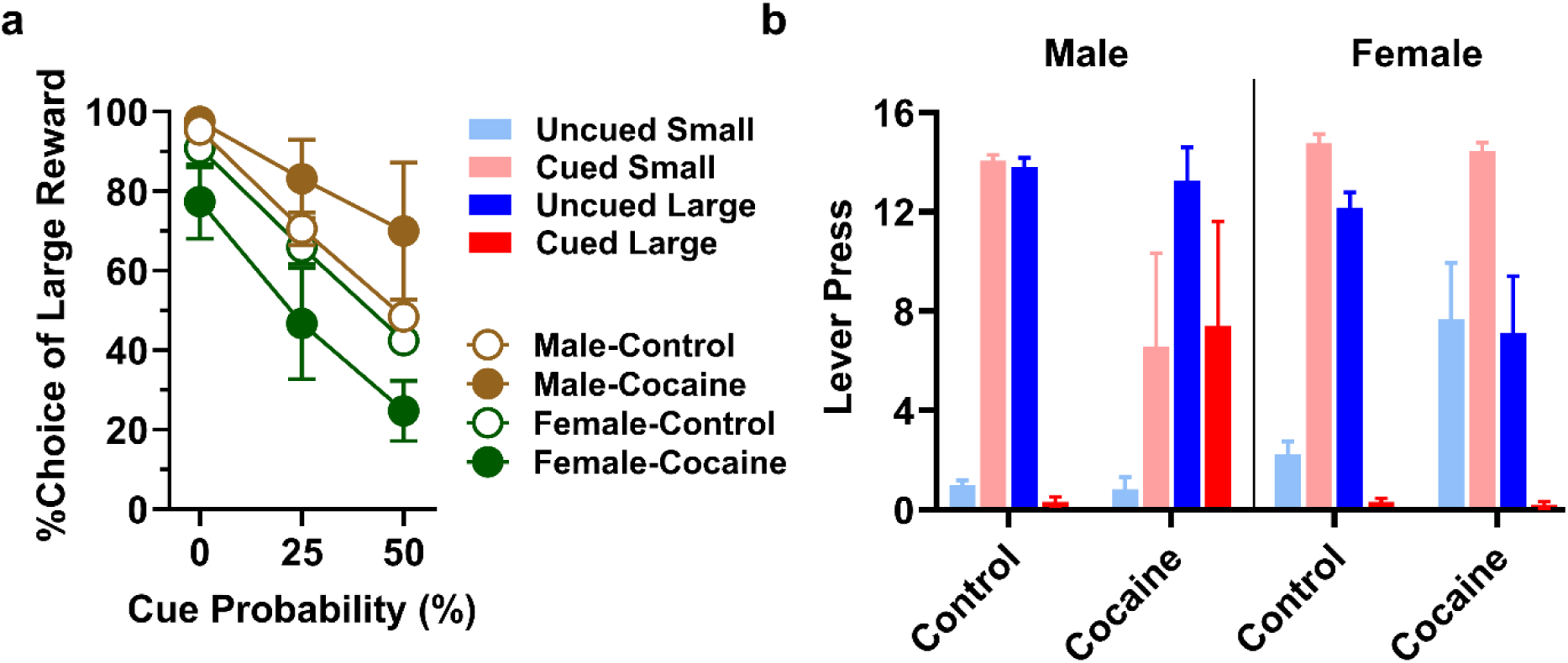
Experiment 2, Effects of chronic cocaine on choice behavior in the CPDT. **a)** %Choice of the large reward decreased as the probability of the punishment cue increased (p<0.01), and male rats chose the large reward more frequently than females (p<0.01). Importantly, in males, prior cocaine increased choice of the large reward compared to controls, whereas in females, prior cocaine decreased choice of the large reward compared to controls (p=0.04). **b)** The presence of the punishment cue decreased the number of large reward choices (p<0.01), and females chose the large reward less frequently than males (p=0.01). Prior cocaine increased the number of large rewards on cued compared to uncued trials (p<0.01), and in males compared to females (p=0.04). Data are represented as means ± SEM.

#### 3.2.2. Choice distribution at 50% probability of punishment cue

Similar to Experiment 1, subsequent analyses focused on data from the two blocks with 50% probability of cue presentation (Figure 4b). A three-factor ANOVA (Sex x Cocaine x Cue) focused on the number of large rewards chosen revealed a main effect of Cue (F(1,14)=88.74, p<0.01) such that rats chose the large reward less frequently on cued trials, and a main effect of Sex (F(1,14)=8.05, p=0.01) such that females chose the large reward less frequently than males. Although there was not a main effect of Cocaine (F(1,14)=0.07, p=0.8), cocaine increased the number of large rewards on cued trials (Cocaine x Cue, F(1,14)=9.62, p<0.01), and in males (Sex x Cocaine, F(1,14)=4.92, p=0.04). No other interaction concerning Sex was significant (Cue x Sex, F(1,14)=0.02, p=0.9; Cue × Cocaine × Sex, F(1,14)=0.46, p=0.51). A similar analysis on the number of small rewards revealed a main effect of Cue (F(1,14)=88.61, p<0.01) such that rats chose the small reward more frequently on cued trials, and a main effect of Sex (F(1,14)=13.88, p<0.01) such that females chose the small reward more frequently than males. Although there was not a main effect of Cocaine (F(1,14)=0.32, p=0.58), cocaine decreased on cued trials and increased on uncued trials the number of small rewards (Cocaine x Cue, F(1,14)=10.33, p<0.01). Cocaine decreased in males and increased in females the number of small rewards (Sex x Cocaine, F(1,14)=8.18, p=0.01). No other interaction concerning Sex was significant (Cue x Sex, F(1,14)=0.01, p=0.91; Cue × Cocaine × Sex, F(1,14)=0.16, p=0.70). These results indicate that punishment cue presentation shifted choices to the small reward and that cocaine amplified this effect in females but damped this effect in males.

Latencies to press levers on free-choice trials at 50% probability of punishment cue were analyzed using a multi-factor repeated-measures ANOVA (Sex x Cue x Cocaine x Lever). The punishment cue reduced latencies to choose levers (F(1,36.55)=8.53, p<0.01), and latencies to choose the small reward were longer than latencies to choose the large reward (Lever, F(1,37.36)=7.83, p<0.01). There was also an interaction between Cue and Lever (F(1,35.20)=5.90, p=0.02), such that latencies to choose the small reward were longer when the punishment cue was absent. In addition, there was an interaction between Sex and Cocaine (F(1,12.93)=20.67, p<0.01), such that Cocaine decreased latencies in females but increased latencies in males, and this effect was greater when choosing the small reward lever (Sex x Cocaine x Lever, F(1,37.36)=5.10, p=0.03). There was no main effect of Cocaine or Sex, nor was there any other interactions concerning these factors (Tables 3 and 4).

**Table 3.**
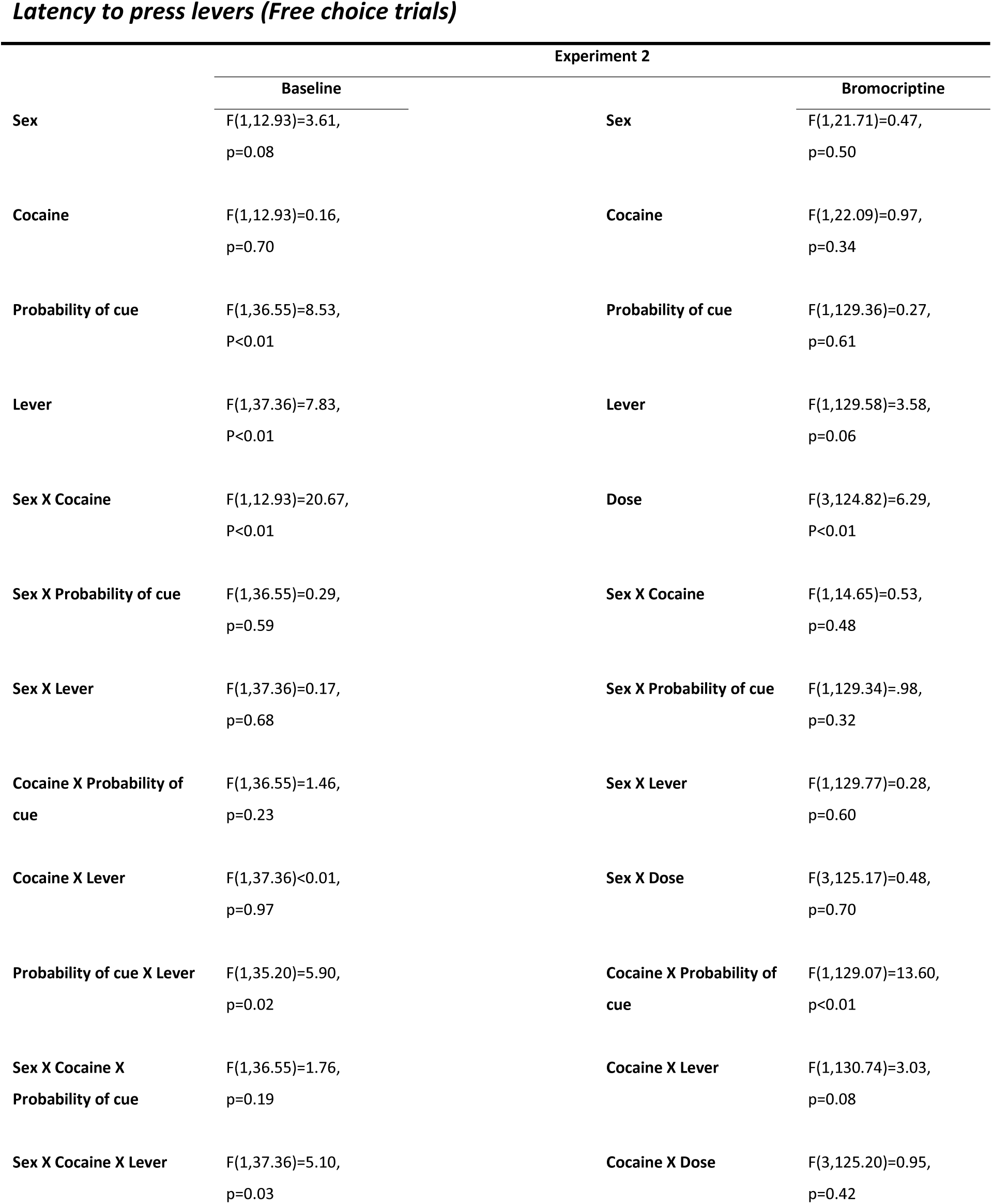
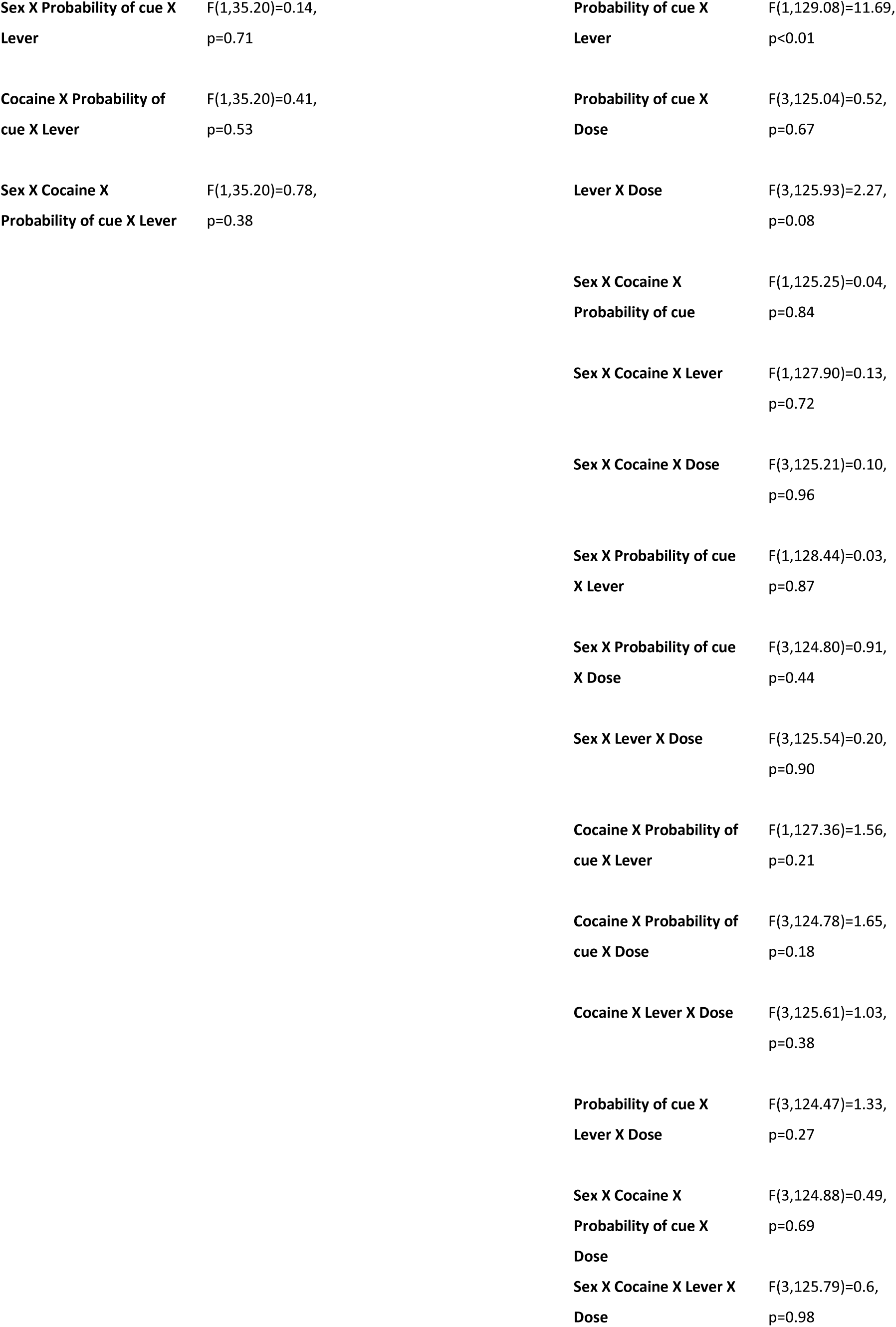
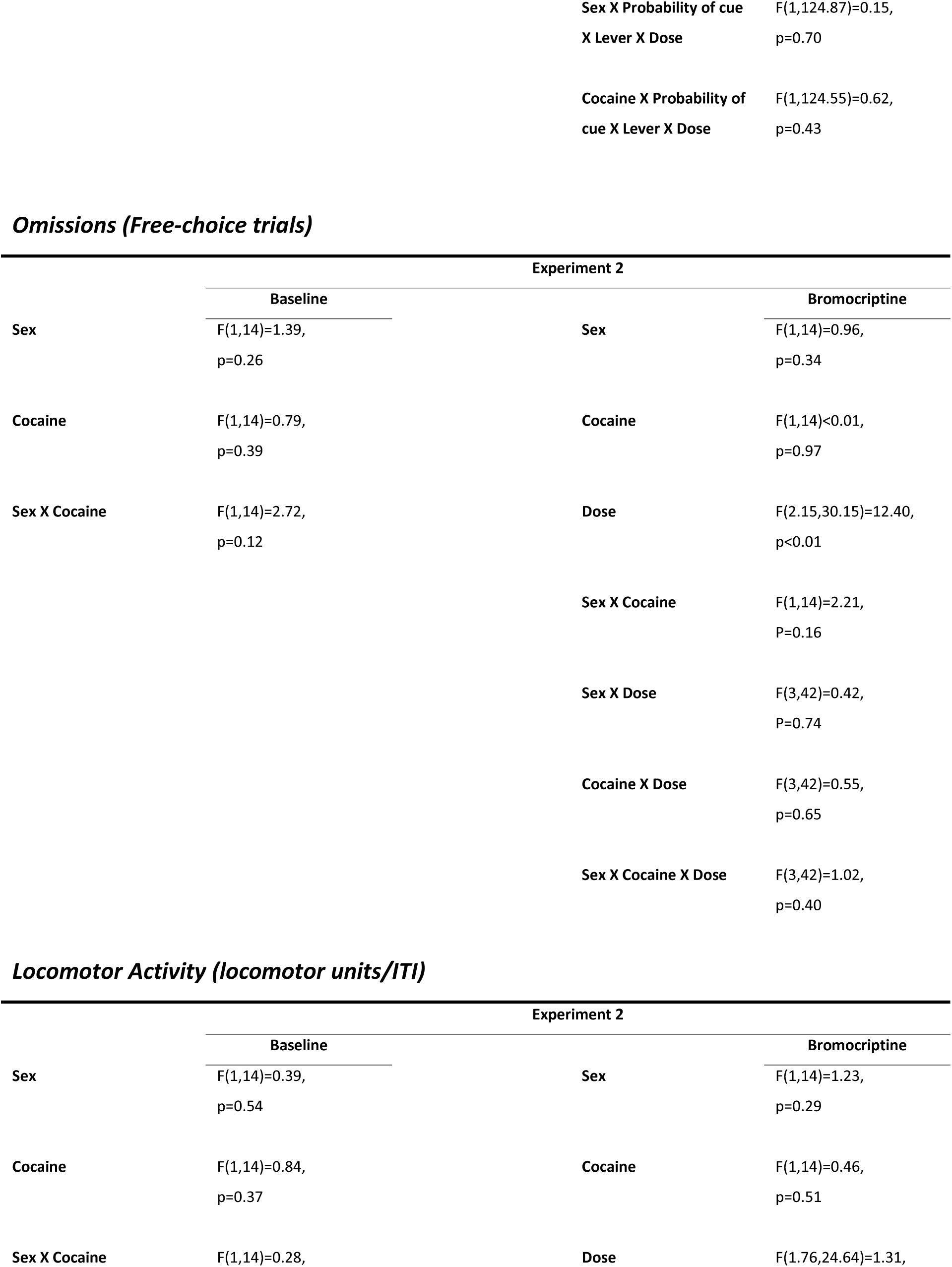
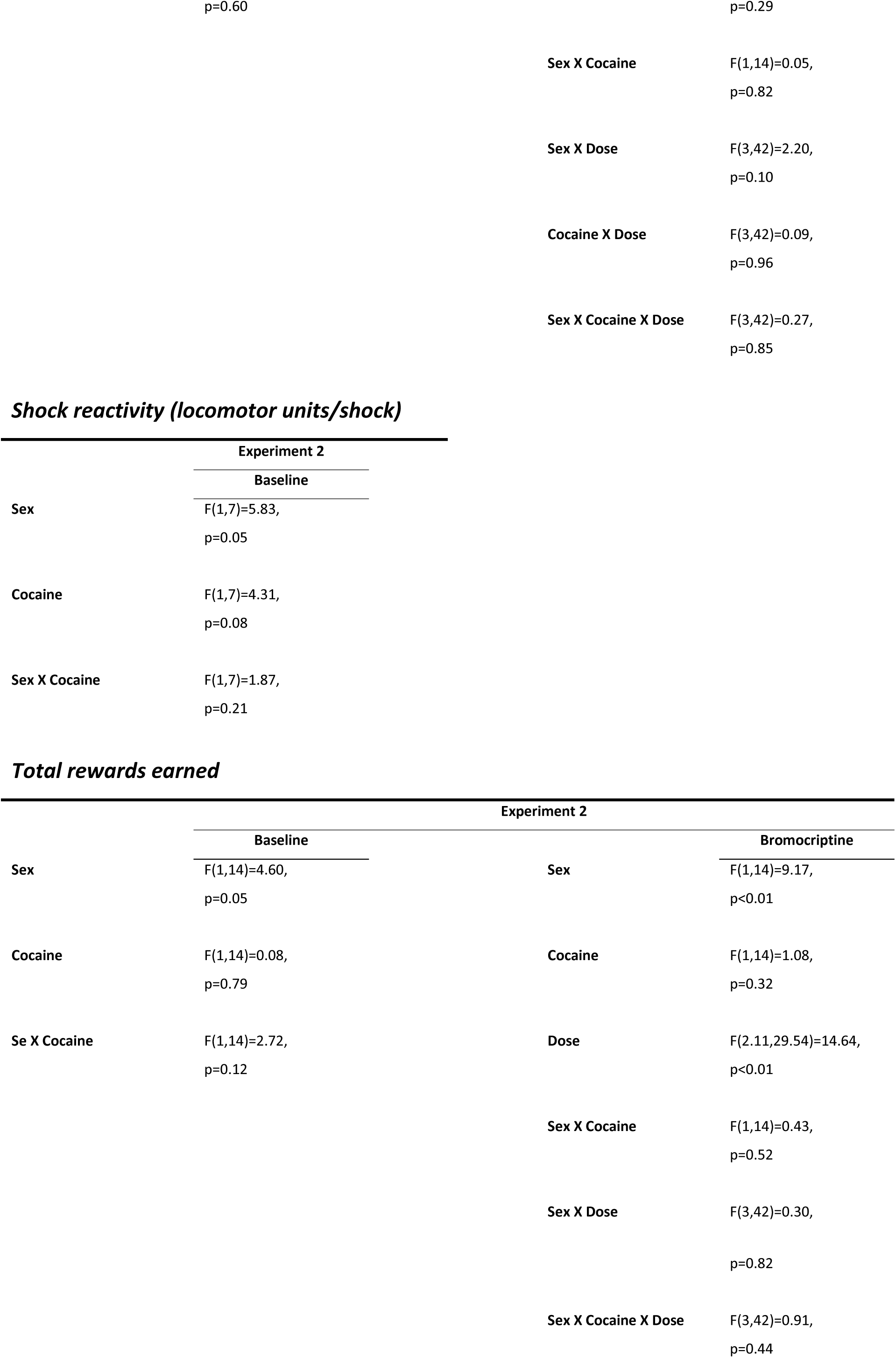
Ancillary measures (statistical results), experiment 2.

**Table 4.** Ancillary measures (mean ± standard error of the mean), experiment 2.

|  |  | *Latency to press levers (Free choice trials)* |  |  |  |
| --- | --- | --- | --- | --- | --- |
|  |  | Cued |  | Uncued |  |
|  |  | Small | Large | Small | Large |
| Control | Female | 0.96 (0.37) | 0.55 (0.24) | 0.92 (0.17) | 0.84 (0.26) |
|  | Male | 0.72 (0.27) | 1.07 (0.74) | 2.06 (0.89) | 0.73 (0.11) |
| Cocaine | Female | 0.39 (0.20) | 1.60 (0.72) | 0.81 (0.26) | 0.69 (0.13) |
|  | Male | 0.59 (1.87) | 0.95 (0.20) | 0.56 (0.12) | 0.77 (0.11) |

|  |  | *Locomotor Activity*<br>*(locomotor units/ITI)* | *Shock reactivity*<br>*(locomotor*<br>*units/shock)* | *Omissions (Free-*<br>*choice trials)* | *Total rewards*<br>*earned* |
| --- | --- | --- | --- | --- | --- |
| Control | Female | 23.97 (16.61) | 3.00 (0.50) | 1.17 (1.38) | 77.50 (2.41) |
|  | Male | 15.79 (8.51) | 1.67 (0.58) | 1.87 (2.81) | 79.67 (2.56) |
| Cocaine | Female | 26.70 (18.84) | 3.25 (1.06) | 1.33 (1.70) | 69.07 (9.44) |
|  | Male | 26.06 (13.91) | 2.88 (0.12) | 5.00 (7.41) | 85.67 (16.23) |

**Bromocriptine**
|  |  |  | Latency to press levers (Free choice trials) |  |  |  |
| --- | --- | --- | --- | --- | --- | --- |
|  |  |  | Cued |  | Uncued |  |
|  |  |  | Small | Large | Small | Large |
| Vehicle | Control | Female | 0.70 (0.24) | 1 | 1.63 (1.27) | 0.72 (0.18) |
|  |  | Male | 0.56 (0.22) | -- | 1.60 (0.97) | 0.53 (0.15) |
|  | Cocaine | Female | 0.40 (0.26) | -- | 1.32 (1.14) | 0.83 (0.20) |
|  |  | Male | 0.30 (0.01) | 0.74 (0.02) | 1.71 (0.66) | 0.74 (0.22) |
| 1.0 mg/kg | Control | Female | 1.86 (0.69) | -- | 2.39 (0.27) | 1.14 (0.48) |
|  |  | Male | 0.97 (0.44) | 4.87 | 1.27 (0.41) | 0.66 (0.16) |
|  | Cocaine | Female | 0.45 (0.28) | 2.11 | 3.03 (3.44) | 1.11 (0.32) |
|  |  | Male | 0.43 (0.16) | 0.91 (0.12) | 2.43 (1.53) | 1.07 (0.44) |
| 3.0 mg/kg | Control | Female | 2.88 (1.25) | -- | 1.97 (0.48) | 0.98 (0.01) |
|  |  | Male | 2.06 (1.65) | 1.29 | 2.84 (1.66) | 0.85 (0.33) |
|  | Cocaine | Female | 1.42 (1.30) | 1.84 | 2.77 (1.14) | 1.47 (0.14) |
|  |  | Male | 2.09 (1.81) | 1.10 | 3.23 (2.17) | 1.17 (0.52) |
| 5.0 mg/kg | Control | Female | 3.13 (3.37) | -- | 1.95 (0.84) | 1.05 (0.39) |
|  |  | Male | 1.84 (1.44) | -- | 2.24 (0.72) | 0.98 (0.24) |
|  | Cocaine | Female | 2.73 (2.04) | -- | 2.82 (1.36) | 1.34 (0.05) |
|  |  | Male | 1.39 (0.94) | 1.09 (0.92) | 3.39 (1.30) | 1.99 (0.94) |

|  |  |  | *Locomotor Activity*<br>*(locomotor units/ITI)* | *Shock reactivity*<br>*(locomotor*<br>*units/shock)* | *Omissions (Free-*<br>*choice trials)* | *Total rewards*<br>*earned* |
| --- | --- | --- | --- | --- | --- | --- |
| Vehicle | Control | Female | 25.97 (18.94) | 1 | 0.00 (0.00) | 79.75 (2.75) |
|  |  | Male | 20.25 (11.02) | -- | 0.20 (0.45) | 77.25 (4.50) |
|  | Cocaine | Female | 24.00 (17.69) | -- | 0.80 (0.45) | 68.20 (9.42) |
|  |  | Male | 28.67 (19.65) | 2.88 (0.00) | 0.75 (0.96) | 86.80 (9.98) |
| 1.0 mg/kg | Control | Female | 22.73 (20.47) | -- | 6.50 (9.71) | 69.00 (17.30) |
|  |  | Male | 13.50 (9.63) | 5 | 1.60 (3.05) | 80.50 (3.42) |
|  | Cocaine | Female | 27.28 (14.64) | 2 | 1.80 (2.17) | 64.80 (15.01) |
|  |  | Male | 20.05 (15.96) | 2.87 (0.09) | 1.25 (0.50) | 82.60 (11.97) |
| 3.0 mg/kg | Control | Female | 34.29 (19.06) | -- | 14.25 (7.14) | 51.25 (14.24) |
|  |  | Male | 14.00 (10.66) | 4 | 5.60 (6.88) | 66.75 (11.95) |
|  | Cocaine | Female | 39.36 (17.86) | 4 | 9.00 (8.28) | 50.00 (9.72) |
|  |  | Male | 22.60 (17.67) | 2.56 | 9.50 (15.84) | 64.80 (28.68) |
| 5.0 mg/kg | Control | Female | 26.46 (17.55) | -- | 12.50 (7.77) | 54.50 (20.04) |
|  |  | Male | 22.14 (25.71) | -- | 4.80 (6.38) | 70.25 (4.35) |
|  | Cocaine | Female | 31.93 (12.20) | -- | 8.60 (6.43) | 42.80 (6.76) |
|  |  | Male | 25.32 (33.10) | 2.83 (0.76) | 13.00 (9.38) | 54.00 (20.02) |

A two-factor ANOVA (Cocaine x Sex) was used to evaluate locomotor activity during the CPDT. There was no main effect of Cocaine or Sex on locomotor activity during the ITI, nor was there an interaction between the factors (Tables 3 and 4). Locomotor activity during shock deliveries in the task was greater in females (Sex, F(1,7)=5.83, p=0.05), but was not affected by Cocaine, nor was there an interaction between Cocaine and Sex (Tables 3 and 4). A two-factor ANOVA (Cocaine x Sex) on the number of omitted trials revealed no main effect of either Cocaine or Sex, nor was there an interaction between the factors (Tables 3 and 4). Analysis of total food rewards earned (Cocaine x Sex) revealed that males earned more rewards than females (F(1,14)=4.60, p=0.05), but Cocaine did not affect earnings, nor was there an interaction between Sex and Cocaine (Tables 3 and 4).

#### 3.2.3. Effects of bromocriptine administration on choice of the large reward

Previous work has shown that the D2/3 dopamine receptor agonist bromocriptine attenuates preference for large, probabilistically-punished rewards [33, 34]. To determine whether this effect extended to conditions in which punishment was explicitly cued, we examined rats’ choice behavior in the CPDT following acute IP administration of bromocriptine (Figure 5). A multi-factor repeated measures ANOVA (Cocaine x Sex x Cue Probability x Bromocriptine) conducted on % choice of the large reward revealed a main effect of Sex (F(1,14)=8.42, p=0.01) such that males chose the large reward more frequently than females, and Cue Probability, such that choice of the large reward declined as the probability of the punishment cue increased (F(1.36,19.01)=59.13, p<0.01). In addition, choice of the large reward decreased following administration of bromocriptine (F(1.97,27.62)=10.73, p<0.01). Cocaine increased choice of the large reward in males, but decreased the same measure in females (Cocaine x Sex, F(1,14)=11.26, p<0.01). The main effect of Cocaine, however, was not significant (F(1,14)=1.80, p=0.20). There were no other significant main effects or interactions concerning Bromocriptine or Cocaine (Cue Probability x Cocaine, F(1.36,19.01)=2.08, p=0.16; Cue Probability x Sex, F(1.36,19.01)=0.89, p=0.39; Bromocriptine x Cocaine, F(1.97,27.62)=0.33, p=0.72; Bromocriptine x Sex, F(1.97,27.62)=0.56, p=0.57; Cue Probability x Bromocriptine, F(3.01,42.13)=1.16, p=0.34; Cue Probability x Cocaine x Sex, F(1.36,19.01)=0.66, p=0.47; Cue Probability x Cocaine x Bromocriptine, F(3.01,42.13)=0.78, p=0.51; Cue Probability x Sex x Bromocriptine, F(3.01,42.13)=1.84, p=0.16; Cocaine x Sex x Bromocriptine, F(1.97,27.62)=0.23, p=0.80; Cue Probability x Cocaine x Sex x Bromocriptine, F(3.01,42.13)=1.44, p=0.24).

**Figure 5.**
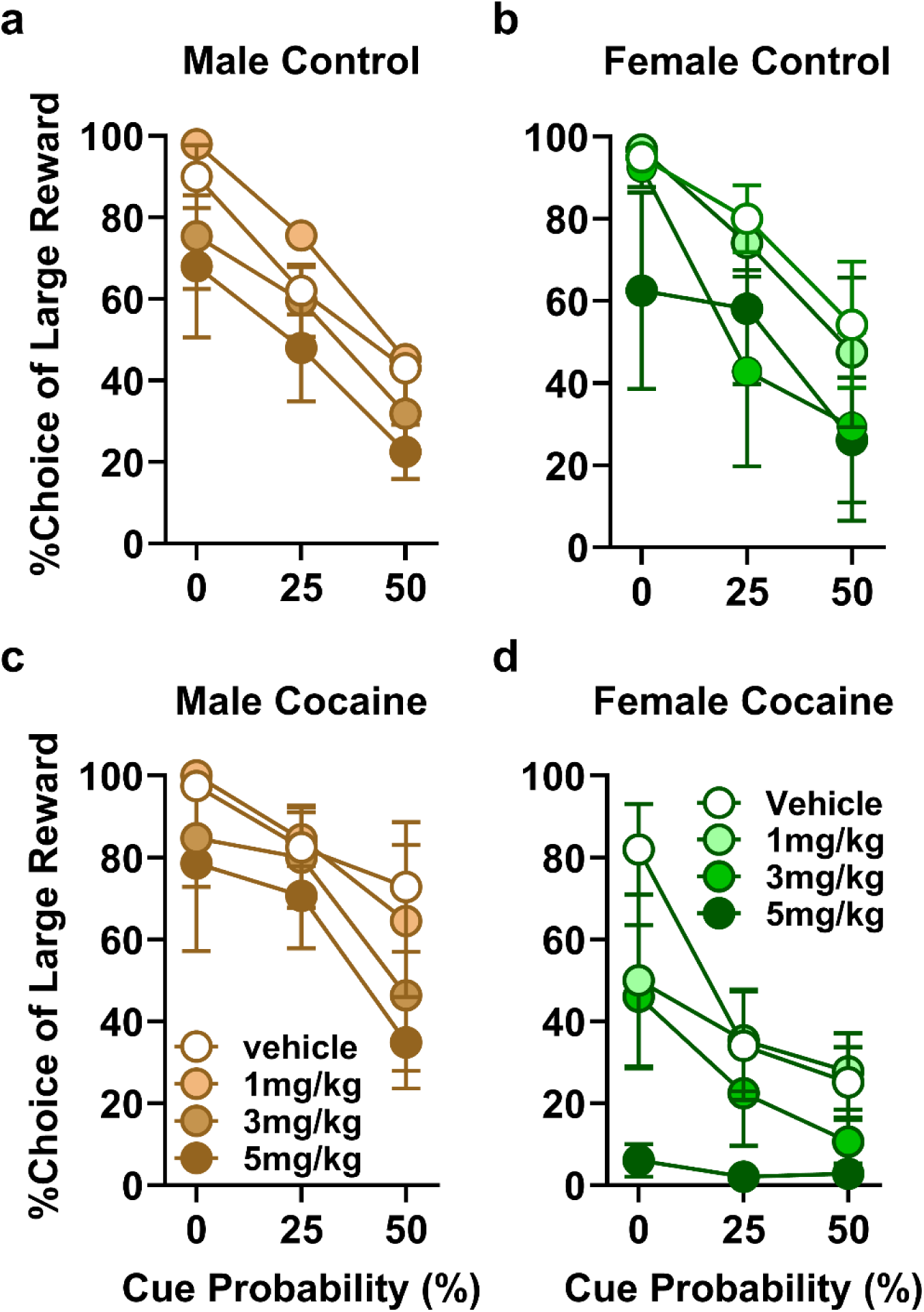
Experiment 2, Effects of acute bromocriptine on % choice of the large reward in rats with a history of chronic cocaine vs. controls. %Choice of the large reward decreased as the probability of the punishment cue increased (p<0.01), and male rats chose the large reward more frequently than females (p<0.01). Bromocriptine decreased %choice of the large reward (p<0.01). Chronic cocaine increased %choice of the large reward in males, but decreased the same measure in females (p<0.01). Data are represented as means ± SEM.

#### 3.2.4. Effects of bromocriptine administration on choice distribution at 50% probability of punishment cue

Subsequent analyses focused on data from the two blocks with 50% probability of cue presentation (Figure 6). A multi-factor ANOVA (Cocaine x Sex x Cue x Bromocriptine) on the number of large rewards revealed a main effect of Cue (F(1,14)=61.83, p<0.01) such that the number of large rewards chosen on cued trials was significantly fewer than on uncued trials. In addition, the number of large rewards decreased following administration of bromocriptine (F(2.12,29.71)=12.03, p<0.01). A Dunnett’s pos-hoc analysis showed that the effect of bromocriptine was significant at 3.0 and 5.0 mg/kg compared to vehicle (p<0.01), and that this effect was more prominent on cued trials at these doses (Cue x Bromocriptine, F(2.49,34.84)=86.65, p<0.01). In addition, the number of large reward choices was greater in males (Sex, F(1,31.56)=10.92, p<0.01), and while cocaine increased this measure in males, females with a history of cocaine showed fewer large reward choices compared to control females (Cocaine x Sex, (F(1,14)=5.39, p=0.04). There was, however, no main effect of Cocaine (F(1,31.56)=0.30, p=0.60). There was no other significant interaction between the factors (Cocaine x Cue, F(1,14)=4.13, p=0.06; Sex x Cue, F(1,14)=0.82, p=0.38; Cocaine x Bromocriptine, F(2.12,29.71)=0.62, p=0.55; Sex x Bromocriptine, F(2.12,29.71)=0.46, p=0.65; Cocaine x Sex x Cue, F(1,14)=0.01, p=0.92; Cocaine x Sex x Bromocriptine, F(2.12,29.71)=0.54, p=0.60; Cocaine x Cue x Bromocriptine, F(2.49,34.84)=0.93, p=0.42; Sex x Cue x Bromocriptine, F(2.49,34.84)=0.97, p=0.41; Cocaine x Sex x Cue x Bromocriptine, F(2.49,34.84)=1.39, p=0.26).

**Figure 6.**
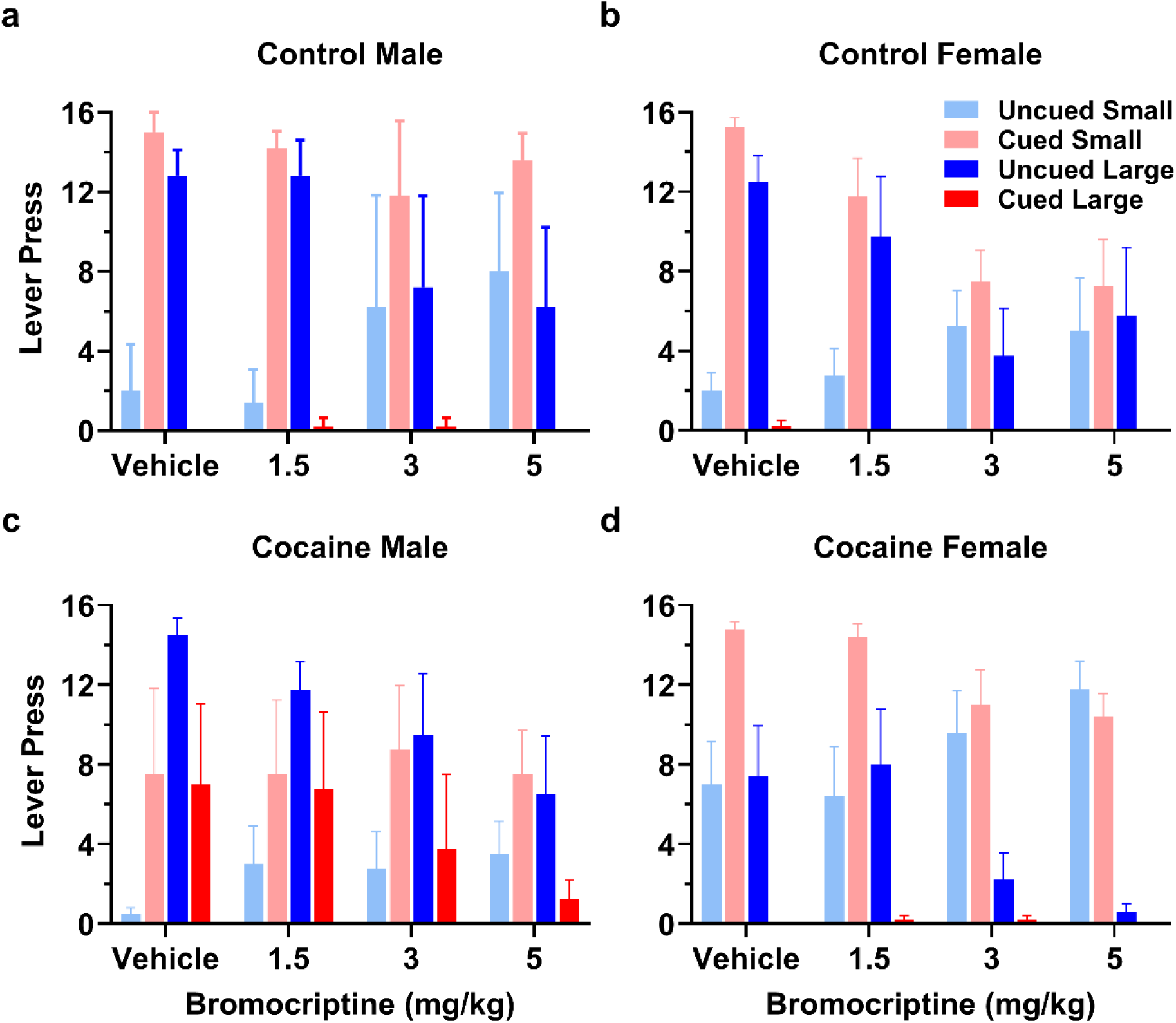
Experiment 2, Effects of acute bromocriptine on choice distribution in rats with a history of chronic cocaine vs. controls. The presence of the punishment cue shifted rats’ choices toward the small reward (p<0.01), and the number of large rewards decreased following administration of bromocriptine (p<0.01). Effects of bromocriptine were more prominent on cued trials (Cue x Bromocriptine, p<0.01). The number of lever presses for large reward was greater in males (p<0.01), but females with a history of cocaine showed fewer lever presses for large rewards (Cocaine x Sex, p=0.04). Data are represented as means ± SEM.

A similar analysis on the number of small reward choices revealed a main effect of Cue (F(1,14)=57.17, p<0.01) such that the number of small reward choices on cued trials was significantly greater than on uncued trials. There was no main effect of bromocriptine (F(2.1,29.40)=0.35, p=0.72); however, bromocriptine decreased the number of small reward choices on cued trials and increased this measure on uncued trials (Cue x Bromocriptine, F(1.94,27.15)=13.81, p<0.01). Sex was not a significant factor in the number of small reward choices (F(1,14)=2.53, p=0.13), nor was Cocaine (F(1,14)=0.20, p=0.89), but Cocaine decreased the number of small rewards in males while increasing the same measure in females (Cocaine x Sex (F(1,14)=10.80, p<0.01). There was no other significant interaction between the factors (Cocaine x Cue, F(1,14)=3.92, p=0.07; Sex x Cue, F(1,14)=1.42, p=0.25; Cocaine x Bromocriptine, F(2.10,29.40)=0.44, p=0.66; Sex x Bromocriptine, F(2.10,29.40)=1.84, p=0.16; Cocaine x Sex x Cue, F(1,14)=0.12, p=0.74; Cocaine x Sex x Bromocriptine, F(2.10,29.40)=0.94, p=0.41; Cocaine x Cue x Bromocriptine, F(1.94,27.15)=1.50, p=0.24; Sex x Cue x Bromocriptine, F(1.94,27.15)=1.64, p=0.21; Cocaine x Sex x Cue x Bromocriptine, F(1.94,27.15)=1.15, p=0.33).

Latencies to press levers on free-choice trials in blocks 3 and 4 were analyzed using a multi-factor repeated measures ANOVA (Sex x Cue x Lever x Cocaine x Bromocriptine). Bromocriptine increased rats’ latencies to press levers (F(3,124.82)=6.29, p<0.01). The punishment cue decreased latencies to choose the small reward lever but increased latencies to choose the large reward lever (Cue x Lever, F(1,129.08)=11.69, p<0.01). In addition, the punishment cue increased latencies to choose levers in the control group but decreased latencies in the cocaine group (Cocaine x Cue (F(1,129.07)=13.60, p<0.01). There was no main effect of either Sex, Cocaine, Cue, or Lever, nor was there any other interactions between factors (Tables 3 and 4).

A three-factor repeated measures ANOVA (Cocaine x Sex x Bromocriptine) was used to evaluate locomotor activity during ITIs in the CPDT, but there were no main effects or interactions involving any of the factors (Tables 3 and 4). Rats omitted the majority of trials when the punishment cue was delivered, and thus, locomotor activity during shock delivery was not analyzed due to the small sample size.

A three-factor repeated measures ANOVA (Cocaine x Sex x Bromocriptine) conducted on the number of omitted trials showed that rats omitted more trials following administration of bromocriptine (F(2.15,30.15)=12.40, p<0.01). A Dunnett’s post-hoc analysis revealed that this effect was significant at the 3.0 and 5.0 mg/kg doses compared to vehicle (p<0.01). There was no main effect of either Cocaine (F(1,14)<0.01, p=0.97) or Sex (F(1,14)=0.96, p=0.34), nor was there an interaction between any of the factors (Tables 3 and 4).

A three-factor repeated measures ANOVA (Cocaine x Sex x Bromocriptine) on the total number of food rewards earned revealed that bromocriptine decreased this measure (F(2.11, 29.54)=14.64, p<0.01), and a Dunnett’s post-hoc analysis showed that this effect was significant at the 3.0 and 5.0 mg/kg doses compared to vehicle. Females earned fewer food rewards than males (Sex, F(1,14)=9.17, p<0.01), but Cocaine did not affect this measure (F(1,14)=1.08, p=0.32). There was no interaction between factors (Tables 3 and 4).

## 4. Discussion

Chronic psychostimulant use can alter behavioral responses to negative outcomes [23, 24, 30, 38–40]. Such altered responses have the potential to precipitate continued substance use and relapse; however, it is less clear how different aspects of such negative outcomes (e.g. intensity, frequency, probability) contribute to these behavioral changes. In previous work, we found that a history of chronic cocaine increases rats’ preference for a large food reward accompanied by probabilistic footshock punishment over a small food reward accompanied by no punishment [24, 41]. Control experiments showed that this cocaine-induced increase in preference for the large, risky reward is not due to reduced sensitivity to footshock [24]. In the present study, we used a novel task (the CPDT) to begin to elucidate whether the increased preference for the probabilistically punished outcome in prior work was due to altered perception of (or preference for) risk vs. altered processing of punishment itself. This task involved a deterministic structure in which the presence of a visual cue prior to each decision between a small and large reward signaled that choice of the large reward would yield a mild footshock alongside the large reward. This design eliminated the risk element associated with choice of the large reward, and enabled investigation of potentially altered processing of the adverse outcome.

The results of Experiment 1 show that in drug-naïve rats of both sexes, the punishment cue was effective in guiding choice behavior. On cued trials, rats preferred the small reward (which was never punished) over the large reward, whereas on uncued trials, rats preferred the large reward over the small reward. Interestingly (and unlike in the RDT, in which females show reduced preference for risky over safe options compared to males [35]), there was no sex difference in the magnitude of this preference. Although this could have been due to the use of different shock intensities in Experiment 1 (lower shock intensities in females than males), the absence of a sex difference in controls in Experiment 2 (in which the same intensity was employed) argues against this interpretation. These data suggest that males and females use punishment cues to guide choice behavior to a comparable extent, and that sex differences observed previously in the RDT are more likely to be driven by differential perception of punishment risk than by differential processing of punishment (or reward) *per se*.

Acute administration of amphetamine at the highest dose (1.5 mg/kg) decreased choice of the large reward compared to vehicle, increased trial omissions, and diminished the efficacy of the punishment cue to guide choice behavior – particularly for choice of the small reward. Previous work in the RDT showed that acute amphetamine decreases preference for large, risky over small, safe rewards in both males and females [34, 35, 42]. Amphetamine can reduce motivation for food intake [43], but this effect is unlikely to explain the reduction in choice of the large reward in either the CPDT or RDT, as in a task similar to the RDT but in which choice of the large reward leads to variable probabilities of reward omission rather than punishment, amphetamine actually increases preference for the large reward [44]. As such, it seems more likely that increased sensitivity to punishment drives the reduced preference for punished over unpunished outcomes. This interpretation is consistent with data showing that acute amphetamine increases the control over instrumental behavior by a conditioned punisher cue [45], and further, that in the CPDT, amphetamine increased the salience of punishment significantly enough to undermine the information delivered by the cue (resulting in decreased choice of the large reward even on uncued trials).

In Experiment 2, control (drug-naïve) males and females did not differ in their choice of the large reward in the CPDT, whereas chronic cocaine increased choice of the large reward in males but decreased the same measure in females. This sex divergence in the effect of chronic cocaine on choice behavior in the CPDT is distinct from our previous findings in the RDT, in which chronic cocaine caused equivalent increases in preference for large, risky rewards in males and females [24]. Notably, however, the current results do suggest that chronic cocaine reduced the extent to which the punishment cue controlled behavior in both sexes, albeit in different ways. Whereas males exposed to cocaine appeared to disregard the cue and continued to choose the large reward when the cue was present, females exposed to cocaine appeared to disregard the absence of the cue, and continued to choose the small reward even when the large reward was indicated as safe. These results are analogous to those from a recent study of the effects of chemogenetic sensitization of ventral tegmental area dopamine neurons on performance of a rat version of the Iowa Gambling Task in the presence or absence of win-paired cues [46]. In this study, sensitization (which mirrored some effects of repeated cocaine) increased the ability of win cues to promote risk taking in males, but reduced the ability of win cues to promote risk taking in females. Considered together, these data could suggest that sensitization of dopaminergic signaling has sex-specific effects on the extent to which cues regulate risk-based decision making, irrespective of whether the cues predict wins or losses, although further research with both sexes using more comparable experimental designs is necessary to address this more conclusively. In addition, the CPDT data suggest that, at least in females, the ability of cocaine to increase preference for large rewards accompanied by risk of punishment in the RDT [24] is not due solely to reduced sensitivity to punishment, as cocaine had directionally-opposite effects in the two tasks.

As an alternative to the idea that chronic cocaine reduced the punishment cue’s control over CPDT choice behavior, the divergent effects on choice behavior in males and females could suggest that the cue provided qualitatively different information to males and females. There is some evidence that in the context of risk-based decision making, males are more sensitive than females to wins/rewards, whereas females are more sensitive than males to losses/punishments [47]. If, despite comparable choice performance under drug-naïve conditions, the absence of the cue (signaling the safety of the large reward) were more salient for males, whereas the presence of the cue (signaling punishment) were more salient for females, then a further, cocaine-induced increase in the salience of these aspects of the cue could have led to the divergent effects on choice behavior observed here.

Systemic administration of the D2/3 dopamine receptor agonist bromocriptine decreased choice of the large reward, increased trial omissions, and reduced sensitivity to the cue in control and cocaine rats of both sexes. These results are similar to those with acute amphetamine in Experiment 1, and consistent with prior work showing that acute bromocriptine has similar effects in the RDT [33, 34]. The neural mechanisms by which bromocriptine and amphetamine exert their effects in the CPDT are unknown. In the RDT, however, acute administration of D2/3 agonists into the nucleus accumbens or basolateral amygdala mimics the effects of systemic bromocriptine [23, 48], suggesting that similar mechanisms could be affected by bromocriptine in the CPDT. Considered together, the results presented here provide a novel context for investigating environmental, behavioral, and neural regulation of cost-benefit decision making, as well as new insights into the mechanisms by which psychostimulants exert lasting effects on cognition and motivated behavior.

## CRediT authorship contribution statement

Mojdeh Faraji: Writing – original draft, Conceptualization, Data curation, Formal analysis. Jennifer Bizon: Writing – review & editing.

Barry Setlow: Writing – review & editing, Conceptualization, Funding acquisition, Supervision.

## Acknowledgement

We thank Vicky Kelley for assistance with behavioral testing, and the Drug Supply Program at the National Institute on Drug Abuse for kindly providing D-amphetamine and cocaine HCl.

Supported by NIH DA036534 (BS).

## Data availability

Data will be made available on request.

## References

1. Gangu, K., et al., Trends of Cocaine Use and Manifestations in Hospitalized Patients: A Cross-Sectional Study. Cureus, 2022. 14(2): p. e22090.

2. Bickel, W.K. and L.A. Marsch, Toward a behavioral economic understanding of drug dependence: delay discounting processes. Addiction, 2001. 96(1): p. 73–86.

3. Richards, J.B., K.E. Sabol, and H. de Wit, Effects of methamphetamine on the adjusting amount procedure, a model of impulsive behavior in rats. Psychopharmacology, 1999. 146(4): p. 432–9.

4. Porter, J.N., et al., Chronic cocaine self-administration in rhesus monkeys: impact on associative learning, cognitive control, and working memory. The Journal of neuroscience : the official journal of the Society for Neuroscience, 2011. 31(13): p. 4926–34.

5. Rogers, R.D., et al., Dissociable deficits in the decision-making cognition of chronic amphetamine abusers, opiate abusers, patients with focal damage to prefrontal cortex, and tryptophan-depleted normal volunteers: evidence for monoaminergic mechanisms. Neuropsychopharmacology : official publication of the American College of Neuropsychopharmacology, 1999. 20(4): p. 322–39.

6. Heil, S.H., et al., Delay discounting in currently using and currently abstinent cocaine-dependent outpatients and non-drug-using matched controls. Addict Behav, 2006. 31(7): p. 1290–4.

7. Lucantonio, F., et al., The impact of orbitofrontal dysfunction on cocaine addiction. Nature neuroscience, 2012. 15(3): p. 358–66.

8. Perry, J.L. and M.E. Carroll, The role of impulsive behavior in drug abuse. Psychopharmacology, 2008. 200(1): p. 1–26.

9. Di Sclafani, V., et al., Neuropsychological performance of individuals dependent on crack-cocaine, or crack-cocaine and alcohol, at 6 weeks and 6 months of abstinence. Drug and alcohol dependence, 2002. 66(2): p. 161–71.

10. Kirby, K.N. and N.M. Petry, Heroin and cocaine abusers have higher discount rates for delayed rewards than alcoholics or non-drug-using controls. Addiction, 2004. 99(4): p. 461–71.

11. Fernandez-Serrano, M.J., et al., Neuropsychological profiling of impulsivity and compulsivity in cocaine dependent individuals. Psychopharmacology, 2012. 219(2): p. 673–83.

12. Schneider, S., et al., Risk taking and the adolescent reward system: a potential common link to substance abuse. The American journal of psychiatry, 2012. 169(1): p. 39–46.

13. Lejuez, C.W., et al., Differences in impulsivity and sexual risk behavior among inner-city crack/cocaine users and heroin users. Drug and alcohol dependence, 2005. 77(2): p. 169–75.

14. Chen, S., et al., Risky decision-making in individuals with substance use disorder: A meta-analysis and meta-regression review. Psychopharmacology, 2020. 237(7): p. 1893–1908.

15. Stevens, L., et al., Disadvantageous Decision-Making as a Predictor of Drop-Out among Cocaine-Dependent Individuals in Long-Term Residential Treatment. Front Psychiatry, 2013. 4: p. 149.

16. Hulka, L.M., et al., Altered social and non-social decision-making in recreational and dependent cocaine users. Psychol Med, 2014. 44(5): p. 1015–28.

17. Bornovalova, M.A., et al., Differences in impulsivity and risk-taking propensity between primary users of crack cocaine and primary users of heroin in a residential substance-use program. Experimental and Clinical Psychopharmacology, 2005. 13(4): p. 311–318.

18. Canavan, S.V., et al., Preliminary evidence for normalization of risk taking by modafinil in chronic cocaine users. Addict Behav, 2014. 39(6): p. 1057–61.

19. Bechara, A., et al., Decision-making deficits, linked to a dysfunctional ventromedial prefrontal cortex, revealed in alcohol and stimulant abusers. Neuropsychologia, 2001. 39(4): p. 376–89.

20. Verdejo-Garcia, A., et al., The differential relationship between cocaine use and marijuana use on decision-making performance over repeat testing with the Iowa Gambling Task. Drug and alcohol dependence, 2007. 90(1): p. 2–11.

21. Simon, N.W., et al., Balancing Risk and Reward: A Rat Model of Risky Decision Making. Neuropsychopharmacology, 2009. 34(10): p. 2208–2217.

22. Orsini, C.A., et al., *Recent Updates in Modeling Risky Decision Making in Rodents*, in Psychiatric Disorders: Methods and Protocols, F.H. Kobeissy, Editor. 2019, Springer New York: New York, NY. p. 79–92.

23. Mitchell, M.R., et al., Adolescent risk taking, cocaine self-administration, and striatal dopamine signaling. Neuropsychopharmacology, 2014. 39(4): p. 955–62.

24. Blaes, S.L., et al., Chronic cocaine causes age-dependent increases in risky choice in both males and females. Behavioral Neuroscience, 2022. 136: p. 243–263.

25. Argyriou, E., et al., Age and impulsive behavior in drug addiction: A review of past research and future directions. Pharmacol Biochem Behav, 2018. 164: p. 106–117.

26. Dolan, S.L., A. Bechara, and P.E. Nathan, Executive dysfunction as a risk marker for substance abuse: The role of impulsive personality traits. Behavioral Sciences & the Law, 2008. 26(6): p. 799–822.

27. Wei, S., et al., Altered Neural Processing of Reward and Punishment in Women With Methamphetamine Use Disorder. Front Psychiatry, 2021. 12: p. 692266.

28. Zhong, N., et al., Smaller Feedback-Related Negativity (FRN) Reflects the Risky Decision-Making Deficits of Methamphetamine Dependent Individuals. Front Psychiatry, 2020. 11: p. 320.

29. Vanderschuren, L.J.M.J. and B.J. Everitt, Drug Seeking Becomes Compulsive After Prolonged Cocaine Self-Administration. Science, 2004. 305(5686): p. 1017-1019.

30. Nguyen, D., et al., Repeated Cocaine Exposure Attenuates the Desire to Actively Avoid: A Novel Active Avoidance Runway Task. Front Behav Neurosci, 2018. 12: p. 108.

31. Jones, B.O., et al., Random interval schedule of reinforcement influences punishment resistance for cocaine in rats. Neurobiology of Learning and Memory, 2024. 213: p. 107961.

32. D’Souza, M.S. and C.L. Duvauchelle, Certain or uncertain cocaine expectations influence accumbens dopamine responses to ‘self-administered cocaine and non-rewarded operant behavior. European Neuropsychopharmacology, 2008. 18(9): p. 628–638.

33. Blaes, S.L., et al., Monoaminergic modulation of decision-making under risk of punishment in a rat model. Behav Pharmacol, 2018. 29(8): p. 745–761.

34. Simon, N.W., et al., Dopaminergic modulation of risky decision-making. J Neurosci, 2011. 31(48): p. 17460–70.

35. Orsini, C.A., et al., Sex differences in a rat model of risky decision making. Behavioral Neuroscience, 2016. 130(1): p. 50–61.

36. Mitchell, M.R., et al., Effects of acute administration of nicotine, amphetamine, diazepam, morphine, and ethanol on risky decision-making in rats. Psychopharmacology (Berl), 2011. 218(4): p. 703–12.

37. Simon, N.W., I.A. Mendez, and B. Setlow, Cocaine exposure causes long-term increases in impulsive choice. Behavioral Neuroscience, 2007. 121(3): p. 543–549.

38. Nguyen, D., et al., Aberrant approach-avoidance conflict resolution following repeated cocaine pre-exposure. Psychopharmacology (Berl), 2015. 232(19): p. 3573–83.

39. Parvaz, M.A., et al., Impaired neural response to negative prediction errors in cocaine addiction. J Neurosci, 2015. 35(5): p. 1872–9.

40. Lim, T.V., et al., Impaired Learning From Negative Feedback in Stimulant Use Disorder: Dopaminergic Modulation. Int J Neuropsychopharmacol, 2021. 24(11): p. 867–878.

41. Orsini, C.A., et al., Distinct relationships between risky decision making and cocaine self-administration under short- and long-access conditions. Prog Neuropsychopharmacol Biol Psychiatry, 2020. 98: p. 109791.

42. Orsini, C.A., et al., Contributions of medial prefrontal cortex to decision making involving risk of punishment. Neuropharmacology, 2018. 139: p. 205–216.

43. Hashemnia, S., D.R. Euston, and A.J. Gruber, Amphetamine reduces reward encoding and stabilizes neural dynamics in rat anterior cingulate cortex. Elife, 2020. 9.

44. St Onge, J.R. and S.B. Floresco, Dopaminergic modulation of risk-based decision making. Neuropsychopharmacology, 2009. 34(3): p. 681–97.

45. Killcross, A.S., B.J. Everitt, and T.W. Robins, Symmetrical effects of amphetamine and alpha-flupenthixol on conditioned punishment and conditioned reinforcement: contrasts with midazolam. Psychopharmacology (Berl), 1997. 129(2): p. 141–52.

46. Hynes, T.J., et al., Win-Paired Cues Modulate the Effect of Dopamine Neuron Sensitization on Decision Making and Cocaine Self-administration: Divergent Effects Across Sex. Biol Psychiatry, 2024. 95(3): p. 220–230.

47. van den Bos, R., J. Homberg, and L. de Visser, A critical review of sex differences in decision-making tasks: focus on the Iowa Gambling Task. Behav Brain Res, 2013. 238: p. 95–108.

48. Wheeler, A.-R., et al., Sex differences in sensitivity to dopamine receptor manipulations of risk-based decision making in rats. Neuropsychopharmacology, 2024. 49(13): p. 1978–1988.

